# Influence of plant genotype and soil on the wheat rhizosphere microbiome: Evidences for a core microbiome across eight African and European soils

**DOI:** 10.1101/777383

**Authors:** Marie Simonin, Cindy Dasilva, Valeria Terzi, Eddy L. M. Ngonkeu, Diégane Diouf, Aboubacry Kane, Gilles Béna, Lionel Moulin

## Abstract

Here, we assessed the relative influence of wheat genotype, agricultural practices (conventional vs organic) and soil type on the rhizosphere microbiome. We characterized the prokaryotic (archaea, bacteria) and eukaryotic (fungi, protists) communities in soils from four different countries (Cameroon, France, Italy, Senegal) and determined if a rhizosphere core microbiome existed across these different countries. The wheat genotype had a limited effect on the rhizosphere microbiome (2% of variance) as the majority of the microbial taxa were consistently associated to multiple wheat genotypes grown in the same soil. Large differences in taxa richness and in community structure were observed between the eight soils studied (57% variance) and the two agricultural practices (10% variance). Despite these differences between soils, we observed that 179 taxa (2 archaea, 104 bacteria, 41 fungi, 32 protists) were consistently detected in the rhizosphere, constituting a core microbiome. In addition to being prevalent, these core taxa were highly abundant and collectively represented 50% of the reads in our dataset. Based on these results, we identify a list of key taxa as future targets of culturomics, metagenomics and wheat synthetic microbiomes. Additionally, we show that protists are an integral part of the wheat holobiont that is currently overlooked.

**Graphical Abstract:** 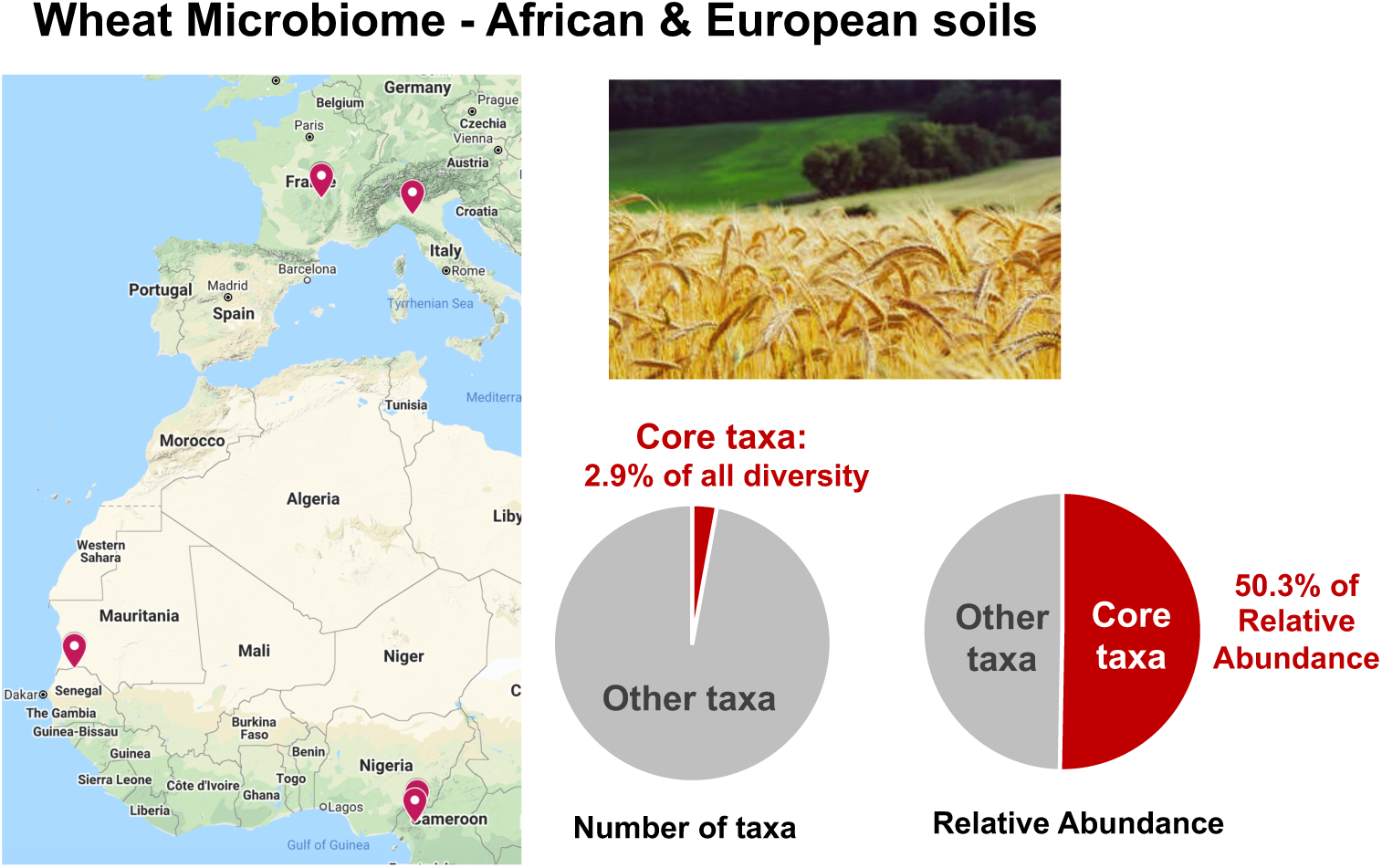

## Introduction

Wheat, with a worldwide production of more than 730 million metric tons in 2018-19 (http://www.fao.org/statistics/en/) is now the second most important grain crop (behind corn) that is mainly used for food, animal fodder, and industrial raw materials. To face the challenges of climate change and of the nutritional needs of a growing world population, there is a necessity of a 70% increase in cereal production by 2050 (FAO 2009). An avenue to improve cereal yields is to harness and manipulate the plant microbiome to improve its nutrition and resistance to pathogens and abiotic stressors (Schlaeppi and Bulgarelli 2015). The plant microbiome is comprised of complex communities of bacteria, archaea, fungi and protists that interact with the host plant in different compartments (e.g. rhizosphere, endosphere, phyllosphere) (Turner, James and Poole 2013). In particular, the root-rhizosphere interface is the nexus of a variety of interactions from which the plant can benefit to acquire mineral nutrients or water and thus is a compartment that is key for future microbiome engineering solutions in agriculture (Bender, Wagg and van der Heijden 2016).

Following the advent of next generation sequencing, several studies characterized the rhizospheric wheat microbiome and investigated the influence of the compartment (rhizoplane vs endosphere), crop management or wheat genotypes on the diversity and structure of these complex microbial communities (Donn *et al.* 2015; Yin *et al.* 2017; Hartman *et al.* 2018; Mavrodi *et al.* 2018). However, a great majority of these studies focused only on bacterial diversity (16S rRNA gene) and more rarely on fungal diversity (ITS) (Sapkota *et al.* 2015; Granzow *et al.* 2017; Lu *et al.* 2018). To our knowledge, no integrative assessment of the wheat rhizospheric microbiome including also the diversity of protists (i.e. amoeba, ciliates, stramenopiles) is currently available. Protists as predators, saprotrophs or phototrophs influence nutrient cycles in the rhizosphere and exert a strong top-down control on microbial (mainly bacterial) biomass and composition (Gao *et al.* 2019). Still, despite protists high diversity and biomass in the rhizosphere, they represent an overlooked component of the plant holobiont.

Moreover, a limited number of studies investigated the effect of the soil used for growing wheat on the rhizosphere microbiome (Fan *et al.* 2017; Mahoney, Yin and Hulbert 2017). For other crops, the culture soil has been demonstrated to be the most important factor structuring the root microbiome before crop management and plant genotype (Lundberg *et al.* 2012; Edwards *et al.* 2015). More work is thus required to study the influence of soil type and geographical location on the wheat microbiome and hence determine which microbial taxa are specific to a soil/location or are common across multiple wheat production systems (i.e. wheat core microbiome). Determining if a wheat core microbiome exists is crucial to help orientate future strain cultivation-based efforts and design microbiome engineering efforts through modifications of agricultural practices or microbial inoculations.

These analyses to determine the presence of a wheat core microbiome need to be conducted at the finest taxonomic level possible (microbial species or strain), as the ecologies and metabolisms of closely related microbial taxa can be extremely different. Previous analyses of wheat microbiome diversity were conducted by lumping amplicon sequences at a 97% identity threshold to create microbial operational taxonomic units (OTUs) that were assumed to represent “species”. New analyses including thousands of genomes indicate that optimal thresholds to represent bacterial species and discriminate phenotypes is at 100% of identity (i.e. exact sequence variant) for the V4 region of the 16S rRNA gene (Edgar 2018). New denoising algorithms for amplicon sequences now enable to eliminate sequencing errors and resolve exact sequence variants (ESVs) that vary only by one nucleotide (Callahan, McMurdie and Holmes 2017). These new bioinformatic methods open new possibilities to characterize plant microbiomes and their thousands of microbial taxa at the finest taxonomic level achievable by short-read amplicon sequencing.

Here, we characterized the rhizosphere microbiome of wheat by considering both prokaryotic (archaea and bacteria) and eukaryotic (fungi and protists) communities of different wheat genotypes grown in soils from four different countries. The goals of this study were to determine the influence of wheat genotype, agricultural practices (conventional vs organic) and soil type on the diversity, structure and taxonomic composition of the rhizosphere microbiome. An additional goal was to determine if a rhizospheric core microbiome existed by identifying microbial taxa present on wheat roots grown in very contrasting soils from different countries. In a growth chamber experiment, we first characterized the rhizosphere microbiome (here soil tightly bound to roots) of eight different genotypes of winter bread wheat (*Triticum aestivum* L.) grown in one soil (FR2) to assess specifically the genotype effect. Second, we characterized the rhizosphere microbiome of three wheat genotypes grown in eight contrasted soils collected from different countries: in Central Africa, Cameroon (CAM1 and CAM2 soils) and West Africa, Senegal (SEN1 and SEN2 soils) and in Europe, France (FR1 and FR2 soils) and Italy (IT1 and IT2 soils). The total microbiome diversity was characterized using amplicon sequencing of the marker genes 16S rRNA (prokaryotic diversity: archaea and bacteria) and 18S rRNA (eukaryotic diversity: fungi and protists). At the taxon level (exact sequence variant), we identified the core microbiome across all soils and genotypes and determined the relative abundance of these core taxa and their potential role as hub taxa in the wheat microbiome using network analyses.

## Methods

### Soil description, sampling and physico-chemical analyses

Eight soils primarily cultivated with wheat which have been previously treated with chemical or organic fertilization were collected from four countries: Senegal and Cameroon (West and Central Africa); France and Italy (Europe). They were chosen for their contrasting physico-chemical and land-use characteristics to represent a diversity of agricultural practices (conventional, organic, agroforestry), texture, pH, carbon and nutrient contents (Table 1). The soils from France and Italy were collected from long-term experimental plots comparing conventional and organic cropping practices and were used to test the influence of agricultural practices on the wheat microbiome. The organic plots received organic fertilization and were managed with a crop rotation (e.g. legume or *Lolium perenne*) while the conventional plots were managed as cereal monocropping with inorganic fertilizers additions. In each field, we collected 12 microsites distant of 20 m from each other and from 0 to 15 cm depth (500 g each). After passing through a 4 mm sieve, samples from the same field were mixed together, a sub-sample was used for physico-chemical analyses, and the remaining was used for pot experiments. The soil samples were shipped to France to the lab IPME to perform the experiments described below. All soil analyses were performed at the Laboratoire des Moyens Analytiques (LAMA) using standard protocols: pH and electrical conductivity (EC) in an aqueous extract (Richards 1954), soil particle size distribution (Bouyoucos 1951), total organic carbon (Pétard 1993), total and available concentrations of nitrogen and phosphorus.

**Table 1.** Main soil characteristics with localization of the collection sites (top) and description of the wheat genotypes used in the study (bottom)

| Soil | Texture | pH | N total (%) | Organic C total (%) | P total (mg/kg) | CaCO<sub>3</sub> total (%) | Agricultural Practices | Lat,Long | Country / region |
| --- | --- | --- | --- | --- | --- | --- | --- | --- | --- |
| CAM1 | Silty-Clay-Loam | 4.5 | 0.92 | 9.3 | 1227 | 0 | Conventional | 5.383333, 10.116667 | Cameroon, Tubah |
| CAM2 | Silt-Loam | 5.7 | 0.90 | 10.2 | 1479 | 0 | Conventional | 5.966667, 10.300000 | Cameroon, Tubah |
| FR1 | Silt-Loam | 8.1 | 0.24 | 2.3 | 1352 | 4.5 | Conventional | 45.777144, 3.142915 | France, Clermont-Ferrand |
| FR2 | Silt-Loam | 8.2 | 0.21 | 4.3 | 1112 | 23.8 | Organic (legume rotation) | 45.768014, 3.157844 | France, Clermont-Ferrand |
| IT1 | Silt | 8.3 | 0.16 | 2.6 | 777 | 12.8 | Conventional | 44.933333, 9.900000 | Italy, Fiorenzuola D'arda PC |
| IT2 | Silt-Loam | 8.1 | 0.26 | 4.4 | 656 | 22.5 | Organic (*Lolium perenne* rotation) | 44.933333, 9.900000 | Italy, Fiorenzuola D'arda PC |
| SEN1 | Clay | 6.3 | 0.12 | 1.5 | 355 | 0 | Conventional | 16.533333, -15.183333 | Senegal, Fanaye (Saint Louis) |
| SEN2 | Loamy Sand | 6.7 | 0.06 | 0.7 | 83 | 0 | Agroforestry | 12.816667, -14.883333 | Senegal, Saré Yéro Bana (Kolda Region) |

| Wheat Genotype | Breeder | Registration Year |
| --- | --- | --- |
| **Apache** | Limagrain, France | 1998 |
| **Bermude** | Florimond Desprez, France | 2007 |
| **Carstens** | Carsten, Germany | 1949 |
| **Champlein** | Benoist, France | 1959 |
| **Cheyenne** | Nebraska Agricultural Experiment Station, USA | 1933 |
| **Rubisko** | RAGT, France | 2012 |
| **Soissons** | Florimond Desprez, France | 1988 |
| **Terminillo** | N.Strampelli, Italy | 1907 |

### Experimental design: 2 sub-experiments

During the same experiment, we performed two sub-experiments in a growth-chamber under controlled conditions in Montpellier (France) with the eight soils collected from African and European countries. The sub-experiment 1 looked at the effects of tender wheat genotype alone on the rhizosphere microbiome by growing eight genotypes (Table 1) in one soil (FR2). The eight winter bread wheats grown in this sub-experiment were: Apache (AP), Bermude (BM), Champlein (CH), Carstens (CT), Cheyenne (CY), Rubisko (RB), Soissons (SS) and Terminillo (TM). These genotypes were selected to cover a diversity that represent traditional and modern cultivars (registration year ranging from 1907 to 2012) that were bred in different countries (Germany, France, USA, Italy).

The Sub-experiment 2 looked at the interactive effects of wheat genotype and soil on the wheat rhizosphere microbiome. Three genotypes from the eight presented above where selected (AP, BM, RB) and were grown in the eight soils collected in Africa and Europe. These three genotypes were selected because they are the most recent genotypes (in terms of date of registration) and are highly cultivated.

### Plant growth and harvest

Seeds were surface disinfected by washing for 40 min in a 9,6% sodium hypochlorite solution (Hurek *et al.* 1994), rinsed 5 times for 3 min in sterile water, then chlorine traces were removed by washing 3 times for 7 min in 2% (w/v) sodium thiosulfate (Miché and Balandreau 2001) and then rinsed 5 times again for 3 min in sterile water, and left in sterile water for another 45 min. Sterilized seeds were then incubated for germination on sterile agar plates (8 g.L^-1^) for 2 days in the dark at 27°C. Five germinated seeds of each genotype were grown on each sampled soil mixed with 50% of sterile sand (size 0.4 to 1.4 mm) in pots of 400 mL with 3 replicates per condition. All pots were incubated in a growth chamber (Temperature day/night: 22°C/18°C, duration day/night: 16 h/8 h, humidity: 60-65%) for a month corresponding to the 3-4 leaf stage at sampling time. No fertilization was applied during the experiment as our objective was to assess the influence of the soil collected itself on wheat microbiome. The plants were watered 3 times a week and were covered with a transparent plastic bag to avoid cross contaminations, especially during watering. Overall, for the sub-experiment 1, eight varieties of wheat were grown in one soil (8 conditions) in triplicates (24 pots). For the sub-experiment 2, three varieties of wheat were grown in eight soils (24 conditions) in triplicates (72 pots). At the end of the experiment, the three tallest plants from each pot were selected. Their roots were collected and shook to remove non-adhering soil, and placed in sterile falcon conical tubes (50 ml) containing 20 mL of NaCl 0.9% to collect rhizosphere soil (tightly bound to roots). These tubes were vortexed for 5 min, roots were removed from the tubes and the tubes were centrifuged for 30 min at 4000 g (Eppendorf). After centrifugation, the pellet constituting the rhizosphere microbiome was flash frozen in liquid nitrogen and stored at −80°C before DNA extraction. The fresh root mass was measured for each plant sample and these values are presented in Figure S1.

### DNA extraction and high-throughput sequencing

Total DNA was extracted using the PowerSoil Extraction Kit (Mo Bio Laboratories, Carlsbad, CA, USA) according to the manufacturer’s instructions. The quality of the DNA was checked by gel electrophoresis and quantified using NanoDrop spectrophotometer. DNA samples were stored at −80°C. The total DNA samples were used as templates using primers F479 and R888 for V4-V5 region of 16S rRNA amplicons and primers FF390 and FR1 for V7-V8 region of 18S rRNA amplicons (Terrat *et al.* 2015). For 16S rRNA gene fragment, 5 ng of template were used for a 25µl of PCR conducted under the following conditions: 94°C for 2 min, 35 cycles of 30 s at 94°C, 52°C for 30 s and 72°C for 1min, following by 7 min at 72°C. Similarly, 18S rRNA gene fragment was amplified under the following PCR conditions: 94°C for 3 min, 35 cycles of 1 min at 94°C, 52°C for 1 min and 72°C for 1 min, following by 5 min at 72°C. For both, PCR was performed using Phusion High-Fidelity DNA Polymerase (BioLabs). The PCR products were purified using QIAquick PCR Purification Kit (QIAGEN) and quantified using the PicoGreen staining Kit. Next steps were performed by Genome Québec platform: a second PCR of nine cycles was conducted twice for each sample under similar PCR conditions with purified PCR products and 10 base pair multiplex identifiers added to the primers at 5’ position to specifically identify each sample and avoid PCR bias. Finally, PCR products were purified and quantified. The high-throughput sequencing was performed on a MiSeq platform (Illumina, San Diego, CA, USA) by Genome Québec platform. All Illumina sequence data from this study were submitted to the European Nucleotide Archive (ENA) under accession number PRJEB34506.

### Bioinformatic analysis of 16S and 18S rRNA gene sequences

The raw 16S rRNA gene and 18S rRNA gene sequences were processed using Qiime 2 (version 2019.1) (Bolyen *et al.* 2019). After performing a quality screening, we used DADA2 (Callahan *et al.* 2016) to process the raw sequences into exact sequence variants (ESVs). DADA2 resolves biological sequences at the highest resolution (level of single-nucleotide differences) that correspond to the best proxy to identify microbial species (Edgar 2018). The DADA2 workflow performs filtering, dereplication, chimera identification, and the merging of paired-end reads. The taxonomic affiliations were assigned using the SILVA 132 database (Quast *et al.* 2012)(Quast et al. 2012) for the 16S rRNA gene sequences and the Protist Ribosomal Reference database (PR^2^) (Guillou *et al.* 2013) for the 18S rRNA gene sequences. ESVs present in only one sample and with less than 10 observations in the entire dataset were excluded. ESVs affiliated to chloroplasts, mitochondria (16S rRNA gene dataset), to Chloroplastida or to Animalia (18S rRNA gene dataset) were removed in order to keep only microbial ESVs. We used negative controls and the *Decontam* package (Davis *et al.* 2018) in R 3.5.2 (R Core Team 2015) to identify contaminant sequences from reagents or introduced during the manipulation of the samples. Four 16S rRNA gene ESVs and seventeen 18S rRNA gene ESVs were identified as contaminant and removed from the dataset. For the 16S rRNA gene dataset, a total of 5145 ESVs and 1455888 reads were present in the final dataset. This prokaryotic dataset (bacteria and archaea) was rarefied to 1018 sequences per sample and 8 samples had to be excluded due to low read numbers after removing non-microbial sequences. For the 18S rRNA gene dataset, a total of 1409 ESVs and 1281880 reads were present in the final dataset. This eukaryotic dataset (micro-eukaryotes) was rarefied to 1127 sequences per sample and 6 samples had to be excluded due to low read numbers after removing non-microbial sequences. Phylogenetic trees of the 16S and 18S rRNA gene ESVs were prepared in Qiime2 (function *qiime phylogeny align-to-tree-mafft-fasttree*) and their visualizations were performed using iTOL (Letunic and Bork 2016).

### Diversity and core microbiome analyses

We explored the effects of the wheat genotype, agricultural practices and soil on the alpha diversity of the rhizosphere microbiome by calculating the observed ESV richness and Faith’s phylogenetic diversity (R package *picante*) (Kembel *et al.* 2010). The effects on microbiome community structure were investigated based on a Bray-Curtis distance matrix visualized using Non-metric Multi-Dimensional Scaling (NMDS, *metaMDS* function) and associated to a permutational multivariate analysis of variance (*adonis* function, 999 permutations) in the R package *vegan* (Oksanen *et al.* 2007). The core taxa representative of the eight African and European soils used in this study were identified based on a criterion of prevalence in at least 25% of the samples (i.e. presence in a minimum of 16 of the 64 samples) with no criterion related to the relative abundance of the taxa, to consider rare but prevalent microbial taxa (function *core_members* in *microbiome* package) (Lahti, Shetty and Blake 2017). Based on this criterion, a list of core taxa was identified and the number and cumulative relative abundance of these core taxa in each sample were calculated. This prevalence criterion enabled the identification of the wheat core microbiome specific of Africa, of Europe and common to both continents. The taxonomic affiliation of the core taxa was verified and refined using nucleotide basic local alignment search tool (BLASTN) analyses.

We inferred cross-domain interaction networks of the wheat core taxa using the SParse InversE Covariance estimation for Ecological Association Inference (*SPIEC-EASI*, version 1.0.2) package in R (Kurtz *et al.* 2015; Tipton *et al.* 2018). We identified hub taxa that may act as potential keystone taxon in an ecological network, following the approach used in Tipton et al. (2018) (Tipton *et al.* 2018) and Agler et al. (2016) (Agler *et al.* 2016). Hub taxa were identified based on their high connectivity and centrality within the network compare to the other taxa using three node parameters: node degree (number of correlations with other taxa), betwenness centrality and closeness centrality. The node parameters were determined using the R package *igraph* (version 1.2.4) (Csardi and Nepusz 2006) and the visualization of the network was performed with the package *ggnet* (version 0.1) (Tyner, Briatte and Hofmann 2017).

The statistical effects of the wheat genotype, agricultural practices and/or soil (and their interaction, when applicable) on the univariate endpoints (e.g. ESV richness, Faith’s phylogenetic diversity, number of core taxa) were assessed using generalized linear models (*glm* function in package *lme4*) and *post hoc* comparisons were performed using the Tukey method (*warp.emm* function in package *emmeans*). Linear regressions using Spearman correlation test were conducted to investigate relationships between the prokaryotic (16S rRNA gene dataset) and eukaryotic (18S rRNA gene dataset) ESV richness or between the number of core taxa and the relative abundance of core taxa per sample. Heatmaps were performed using the R package *pheatmap* and scatter plots were prepared using the package *ggplot2* (version 3.1.1) (Wickham 2016).

### Availability of data and materials

The raw amplicon sequencing data are available on the European Nuclotide Archive (ENA) with the accession number PRJEB34506: http://www.ebi.ac.uk/ena/data/view/PRJEB34506. The code, metadata and datasets used for the bioinformatic analyses to process the amplicon sequencing data and for generating the figures in R are available at the following link: https://github.com/marie-simonin/Wheat_Microbiome.

## Results

### Limited effects of wheat genotype on the rhizosphere microbiome in one soil

We observed limited effects of the wheat genotype on the rhizosphere microbiome after 30 days of growth in experimental chambers under controlled conditions. There were no statistical differences on the total ESV richness (272 to 368 total ESVs on average) and phylogenetic diversity (15.2 to 18.7 total PD) between 7 genotypes for the total microbial diversity (Fig. 1A, 1B). Only the genotype BM presented a lower diversity than some genotypes. Interestingly the genotypes recruited different proportions of prokaryotes and eukaryotes in their rhizospheres. Some genotypes with the highest prokaryotic diversity (AP, RP, TM) presented a lower eukaryotic diversity, while the genotypes that presented the highest eukaryotic diversity (CT, CH, CY) had low to intermediate levels of prokaryotic diversity (Fig. 1A). As a consequence, no significant correlation was observed between prokaryotic and eukaryotic ESV richness because of the contrasted patterns observed between the different genotypes (Fig. 1C). Additionally, no correlation was observed between the fresh root mass and total ESV richness (Fig. S1C, R^2^=0.005, P=0.73).

**Fig. 1:**
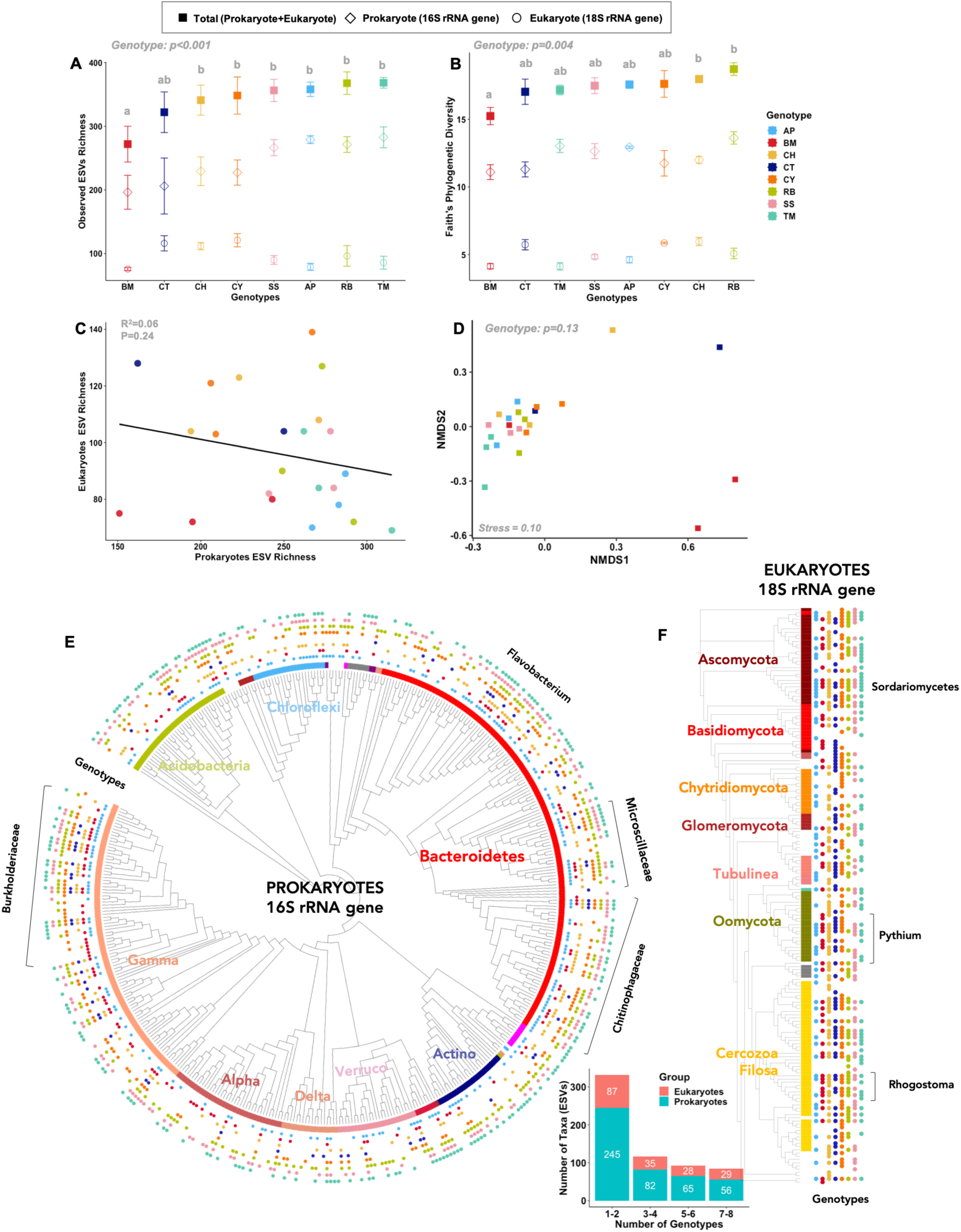
In the soil FR2, influence of 8 wheat genotypes on the rhizosphere microbiome. A) Observed Exact Sequence Variants (ESVs) richness and B) Faith’s phylogenetic diversity of the rhizosphere microbiome for the eight genotypes for the prokaryotic, eukaryotic and total (prokaryotic + eukaryotic) microbial community. Note that on the x axis, the genotypes are ordered from low to high total microbiome diversity. The statistical differences between the eight genotypes are represented by different letters: two genotypes sharing the same letter are not statistically different. For clarity, only the statistical results for the total diversity are presented, detailed results for prokaryotic and eukaryotic diversity are presented in Fig S2. C) Absence of linear correlation between prokaryotic and eukaryotic diversity (ESV richness) across the 8 wheat genotypes. D) Non-metric Multi-Dimensional Scaling (NMDS) ordination showing the absence of significant differences in the structure of the rhizosphere microbiome (prokaryotes and eukaryotes combined) between the eight wheat genotypes. Detailed results for prokaryotic and eukaryotic community structure in Fig S2. Phylogenetic trees of the E) prokaryotic and F) eukaryotic communities showing that many microbial taxa are present in the rhizosphere of multiple wheat genotypes. On the eight outer rings (E) or vertical bands (F), the presence of a colored circle indicates that an ESV was found in the rhizosphere microbiome of a specific wheat genotype. For legibility purposes on the trees and the bar graph, only ESVs that were observed in minimum two replicates of the same genotype condition are represented (n=448 ESVs for prokaryotes and n=179 ESVs for eukaryotes).

No significant effect of wheat genotype was either observed on total microbiome structure (Fig. 1D, P=0.13). This limited genotype effect was also observed when analyzing the patterns separately on prokaryotic (Fig. S2, P=0.04) and eukaryotic community structure (Fig. S2, P=0.22). The phylogenetic trees show that hundreds of microbial taxa are associated to the rhizosphere of multiple wheat genotypes. We observed that 117 microbial taxa were reliably associated (i.e. present in multiple replicates) to 3 or 4 wheat genotypes, 93 taxa were associated to 5-6 genotypes and 85 taxa to 7-8 genotypes (Fig. 1 bar graph). These taxa associated to multiple genotypes were especially affiliated to the bacterial families Burkholderiaceae, Chitinophagaceae, Caulobacteraceae or fungal Ascomycota and cercozoan Filosa for the eukaryotes (list of 85 taxa associated to 7-8 genotypes is available in Additional file 1). Additionally, we observed that the bacterial phyla or class Actinobacteria, Deltaproteobacteria and Alphaproteobacteria were more specifically associated to the genotypes with a high bacterial richness (AP, RP, TM, Fig. 1).

## Characterization of wheat rhizospheric microbiome across eight African and European soils

### Strong effect of soil on the structure and diversity of the rhizosphere microbiome

Cultivation of wheat in different soils led to large differences in the ESV richness (100 to 380 total ESVs) and phylogenetic diversity (8 to 22 total PD) of the rhizosphere microbiome (Fig. 2A, 2B). In most soils, the wheat genotype had no effect on alpha diversity (Fig S3), hence the alpha diversity results in Figure 2 are presented with the data from the three genotypes combined. SEN1 soil had the lowest diversity and SEN2, FR1 and IT2 presented the highest microbiome diversity. The two Cameroonian soils (CAM1 and CAM2) and the FR2 soil presented intermediate levels of diversity (300-350 ESVs). The proportion of prokaryotes and eukaryotes varied between soils but across all soils a positive relationship between prokaryote and eukaryote diversity was found (Fig. 2C, R^2^=0.4, P<0.001). No correlation was observed between the fresh root mass and total ESV richness (Fig. S1D, R^2^=0.03, P=0.22).

**Fig. 2:**
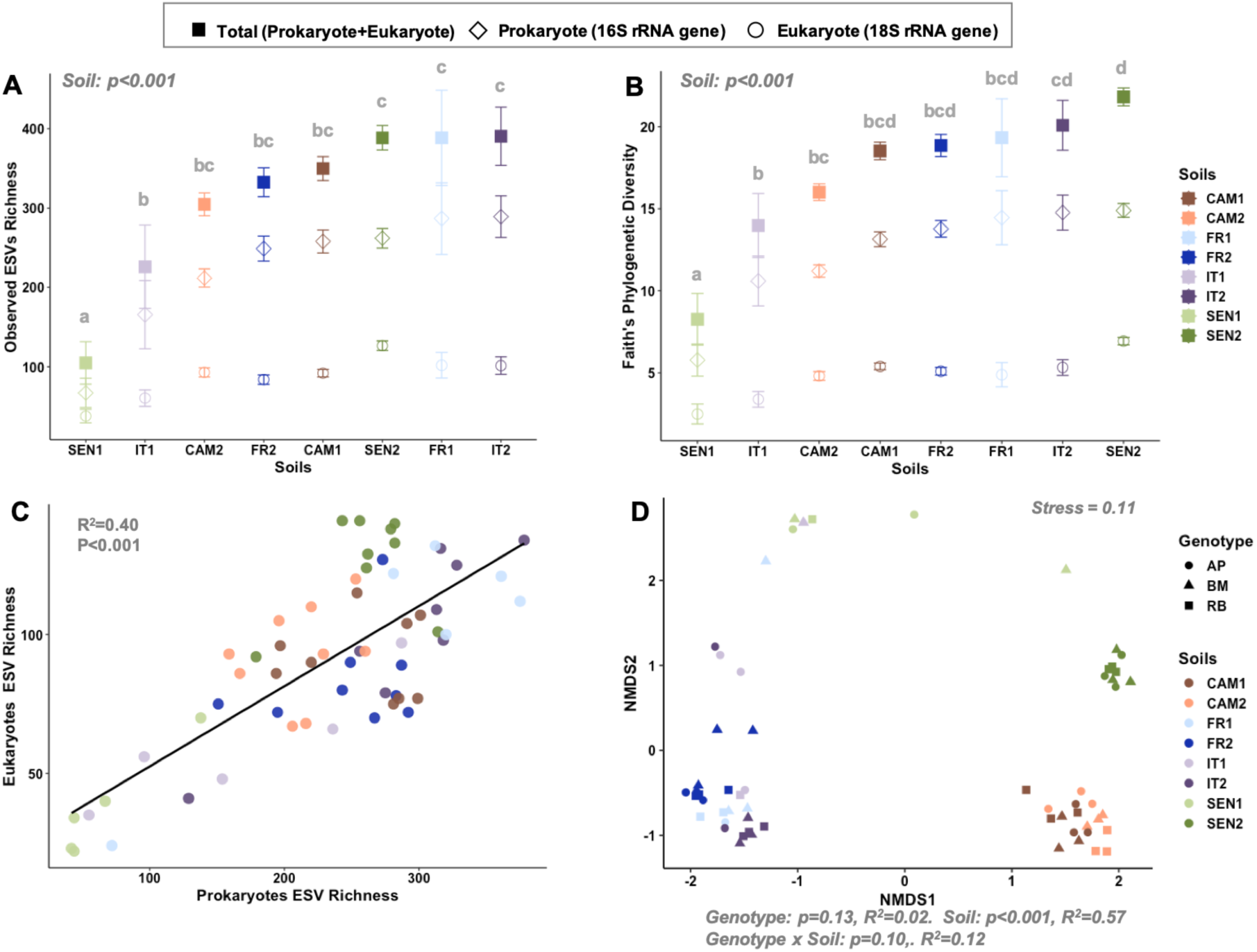
Comparisons of the rhizosphere microbiome across eight agricultural soils from Cameroon (CAM1, CAM2), France (FR1, FR2), Italy (IT1, IT2) and Senegal (SEN1, SEN2). A) Observed Exact Sequence Variants (ESVs) richness and B) Faith’s phylogenetic diversity of the rhizosphere microbiome in the eight soils for the prokaryotic, eukaryotic and total (prokaryotic + eukaryotic) community. Note that on the x axis, the soils are ordered from low to high total microbiome diversity. For the Observed ESV Richness and Faith’s Phylogenetic diversity, the average of the three genotypes is presented for each soil because in most soils the genotype had no effect on alpha diversity (Fig S3). The statistical differences between the eight soils are represented by different letters: two soils sharing the same letter are not statistically different. For clarity, only the statistical results for the total diversity are presented, detailed results for prokaryotic and eukaryotic diversity are presented in Fig S3 and S4. C) Significant positive linear correlation between prokaryotic and eukaryotic diversity (ESV richness) across the eight soils. D) NMDS ordination showing a significant soil effect on the structure of the rhizosphere microbiome (prokaryotes and eukaryotes combined). Detailed results for prokaryotic and eukaryotic community structure in Fig S4.

A very strong effect of the soil was also observed on the structure of the microbial community explaining 57% of the variance, while the effects of wheat genotype (3 genotypes tested, 2% of the variance) and the interactive effects between genotype and soil were not significant (Fig. 2D). The rhizosphere microbiome samples separated along the NMDS axis 1, mainly according to the continent where the soil was collected (African soils on the right and European soils on the left). A clear separation between the Cameroonian and Senegalese soils was also observed on the NMDS axis 2, while the differences in community structure between the two European soils were not as strong.

### Agricultural practices affect the structure and diversity of the rhizosphere microbiome

To assess the influence of agricultural practices, we took advantage of the fact that the Italian and French soils were collected in long-term experimental plots under two types of farming practices at the same location (conventional farming with chemical fertilization, and organic farming with culture rotation with legumes or gramineous plants, Table 1). We observed a significant difference in total taxa richness between the two agricultural practices, only for the Italian soils (Fig. 3A). Taxa richness was 56% higher in the organic Italian soil than in the conventional Italian soil. However, we observed a significant difference in microbiome structure between conventional and organic farming in both Italian and French soils (Fig. 3B). The separations between Conventional and Organic microbiome samples were present on the NMDS axis 2 and the agricultural practices explained globally 10% of the variance in community structure (Fig. 3B). Using differential abundance testing, we identified 51 taxa that were significantly enriched in conventional farming (in Italian or French soils) and 16 taxa in Organic farming (Fig. S5). For the taxa enriched in Conventional farming, half were bacteria and the other half eukaryotes (Cercozoans, Fungi and Oomycetes) including taxa affiliated to *Chitinophaga, Flavobacterium, Bacillus, Pseudomonas, Glissomonadida, Cordyceps* or *Phytophtora*. For the taxa enriched in organic farming, most bacteria were affiliated to *Flavobacterium* and *Cellvibrio*, while the eukaryotes were two uncultured fungi, two uncultured Stramenopiles and the oomycetes *Pythium*.

**Fig. 3:**
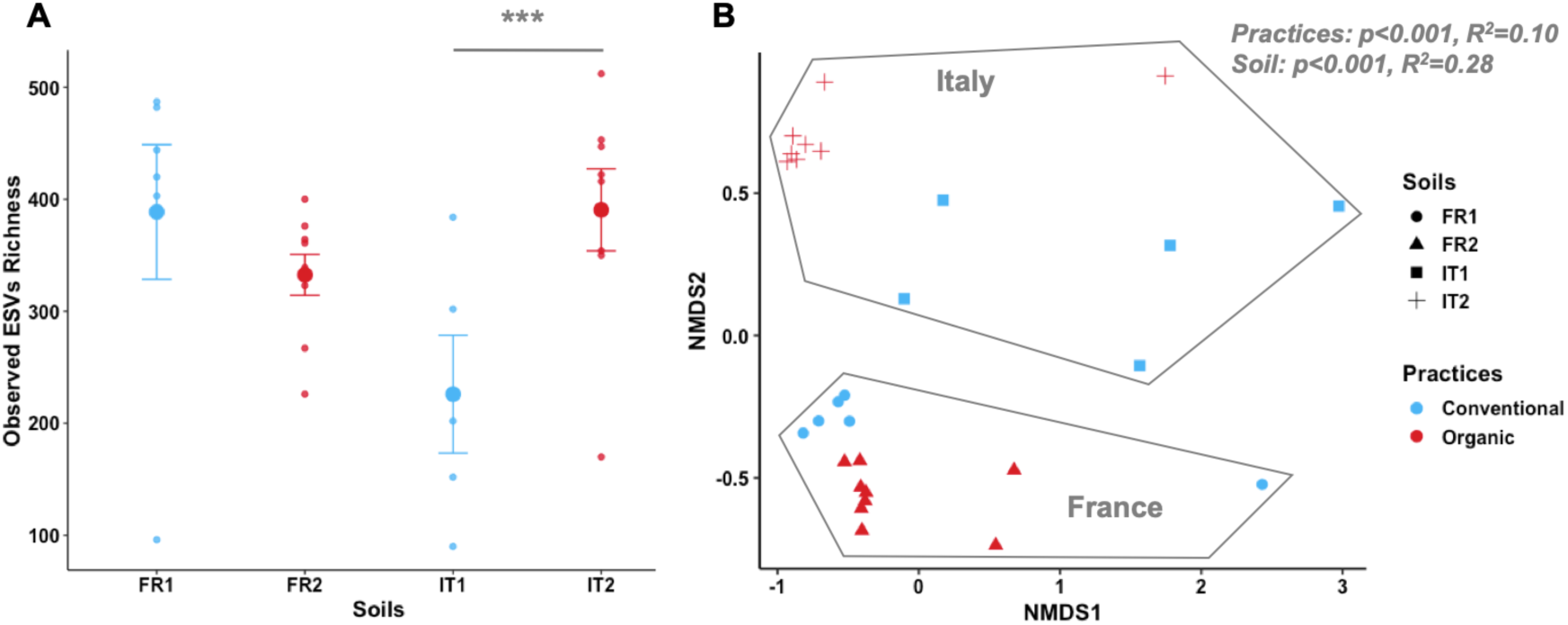
Effect of agricultural practices (conventional vs organic farming) on the rhizosphere microbiome diversity and structure in the French and Italian soils. A) Observed Exact Sequence Variants (ESVs) richness of the rhizosphere microbiome for the prokaryotic, eukaryotic and total (bacterial + eukaryotic) community. B) NMDS ordination showing a significant effect of agricultural practices on the structure of the rhizosphere microbiome (prokaryotes and eukaryotes). The list of the 67 taxa significantly affected by farming practices is available in Additional file 1.

### Phylogenetic distribution of rhizosphere microbial taxa across the eight soils

Across the eight soils studied, we observed that the wheat rhizosphere microbiome was extremely diverse with a total of 4760 ESVs identified. For the Prokaryotes, Archaea represented only 0.9% of the reads (29 ESVs, Phylum Thaumarcheota only) that were affiliated to two archaeal families of ammonia oxidizers (Nitrosophaeraceae and Nitrosotaleaceae) and the remaining 99.1% of the reads belonged to 3652 bacterial ESVs in the final dataset. Across all soils, the most abundant and diverse bacterial families were Burkholderiaceae (10% relative abundance, 251 ESVs, phylum Proteobacteria) followed by Chitinophagaceae (4.2% relative abundance, 296 ESVs, phylum Bacteroidetes) and Flavobacteriaceae (4% relative abundance, 79 ESVs, phylum Bacteroidetes) (bubble plot, Fig. 4). For the Eukaryotes (1125 ESVs), fungal taxa were the most abundant and diverse (61% relative abundance, 459 ESVs), followed by Cercozoa (17%, 415 ESVs) and Oomycota taxa (14%, 47 ESVs). The eukaryotic dominant clades were Sordariomycetes (7.3%, 90 ESVs, Ascomycota), Peronosporales (8.6%, 36 ESVs, Oomycota) and Chytridiomycetes (6.5%, 110 ESVs, Chytridiomycota) (bubble plot, Fig. 4). As seen on the phylogenetic trees, these abundant and diverse prokaryotic or eukaryotic taxa were present in multiple soils (Fig. 4). More specifically, when considering all the prokaryotic and eukaryotic taxa associated to the wheat rhizosphere (n=4760 EVs), we found that 402 taxa (8.5%) where observed in both African (Cameroon and Senegal) and European (France and Italy) soils and that more than 2000 taxa where specific of the 2 continents (Venn diagram, Fig. 4). The phyla in which few taxa were shared between continents (clear continental phylogenetic signals) were principally in the Acidobacteria, Chloroflexi, and Bacteroidetes phyla for Bacteria and the phylogenetic signals were not as clear for eukaryotes (Fig. 4). However, it is important to note that the large majority of prokaryotic and eukaryotic ESVs were observed in multiple countries (Fig. 4).

**Fig. 4:**
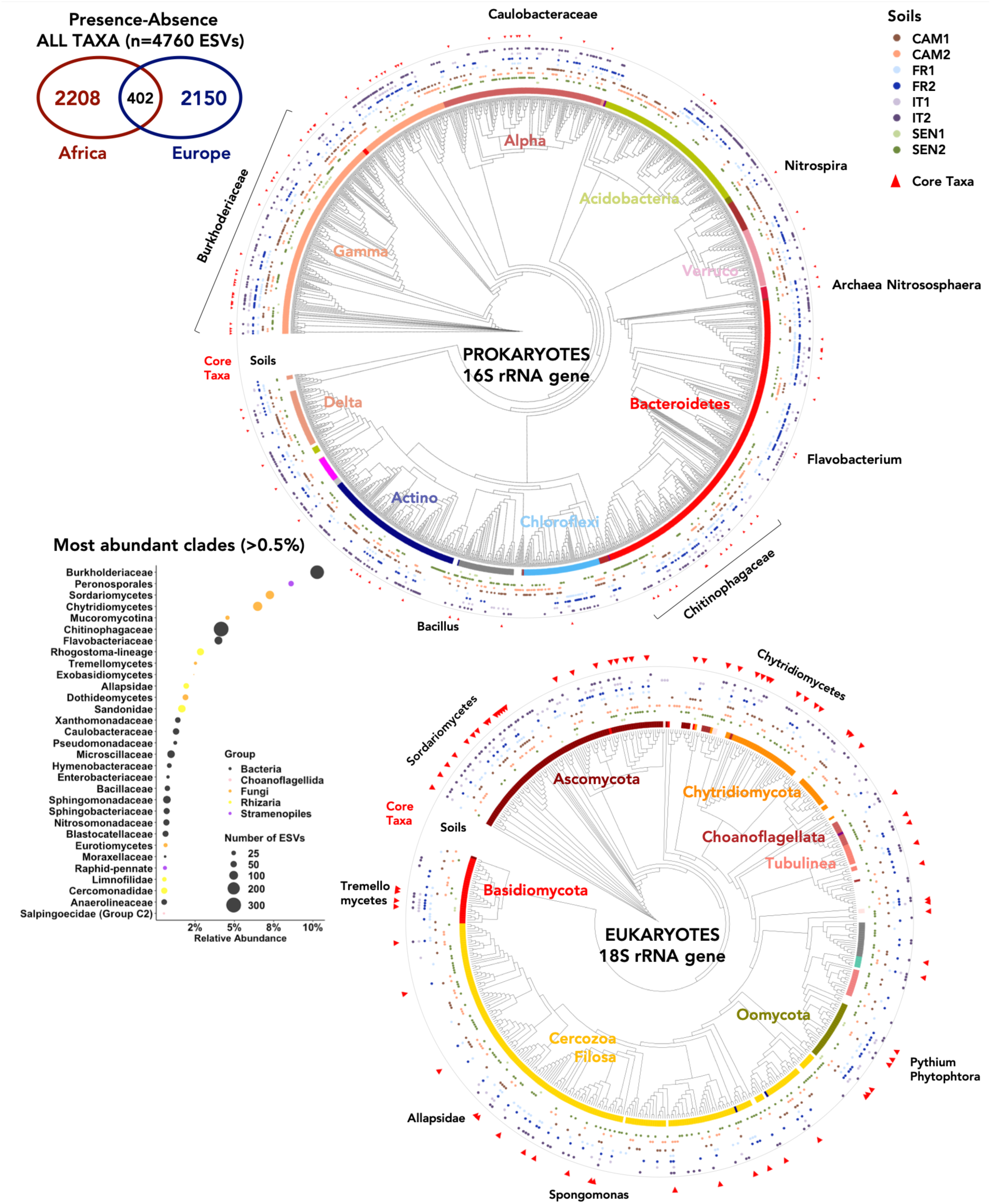
Phylogenetic trees of the prokaryotic (top) and eukaryotic (bottom) taxa present in the eight African and European soils. On the eight outer rings represent the eight soils and the presence of a colored circle indicates that an ESV was found in this soil. Core taxa are indicated by a red triangle on the outer ring. For legibility purposes on the trees, only ESVs that were observed in minimum two replicates of the same genotype-soil condition are represented (n=1478 ESVs for prokaryotes and n=537 ESVs for eukaryotes). In the top-left corner, a Venn diagram represent the number of taxa specific of African or European soils and the number taxa observed in both continents. A bubble plot presenting the relative abundance of the most abundant clades (relative abundance >0.5%) across all samples and the size of the points represent the number of taxa (ESVs) in each clade.

## Evidences for a rhizosphere core microbiome shared across African and European soils

### 2.9% of the microbiome diversity represent 50% of the rhizosphere microbiome abundance

Core taxa (ESV level) of the wheat rhizosphere microbiome were identified based on a criterion of presence of the ESV in 25% of the samples with no relative abundance threshold (core taxa can be rare). This prevalence criterion enabled the identification of the wheat core microbiome specific of Africa, of Europe and common to both continents. The 179 ESVs that responded to the criterion defining a core taxon in our study are indicated by a red triangle on the phylogenetic trees in Fig. 4. These core taxa were distributed across all domains (2 Archaea, 104 Bacteria, 73 Eukaryotes) and spanned 65 families (Fig. 5A, B) and 84 genera. We observed that 118 taxa out of the 179 taxa where observed in soils from both continents (66%), while only 20 core taxa where specific of African soils and 41 of European soils (Fig. 5A). Surprisingly, this list of only 179 ESVs on the total 4760 ESVs (prokaryote + eukaryote) represented 51% of all the sequences in the dataset (Fig. 5D, E) indicating that collectively these 179 core taxa represent half of the relative abundance of the rhizosphere microbiome. From one soil to another, the number of observed taxa belonging to this list of core taxa varied (Fig. 5C) with the SEN1 soil presenting only 22 core taxa on average (total relative abundance of 52%, Fig. 5D) and the FR2 soil presenting 90 core taxa on average (total relative abundance of 65%, Fig. 5D). Interestingly, we did not observe a positive correlation between the number of core taxa observed and their relative abundance in a sample, indicating that across very different soils, these taxa always represented a high relative abundance even when a low number of them were present (Fig. S6).

**Fig. 5:**
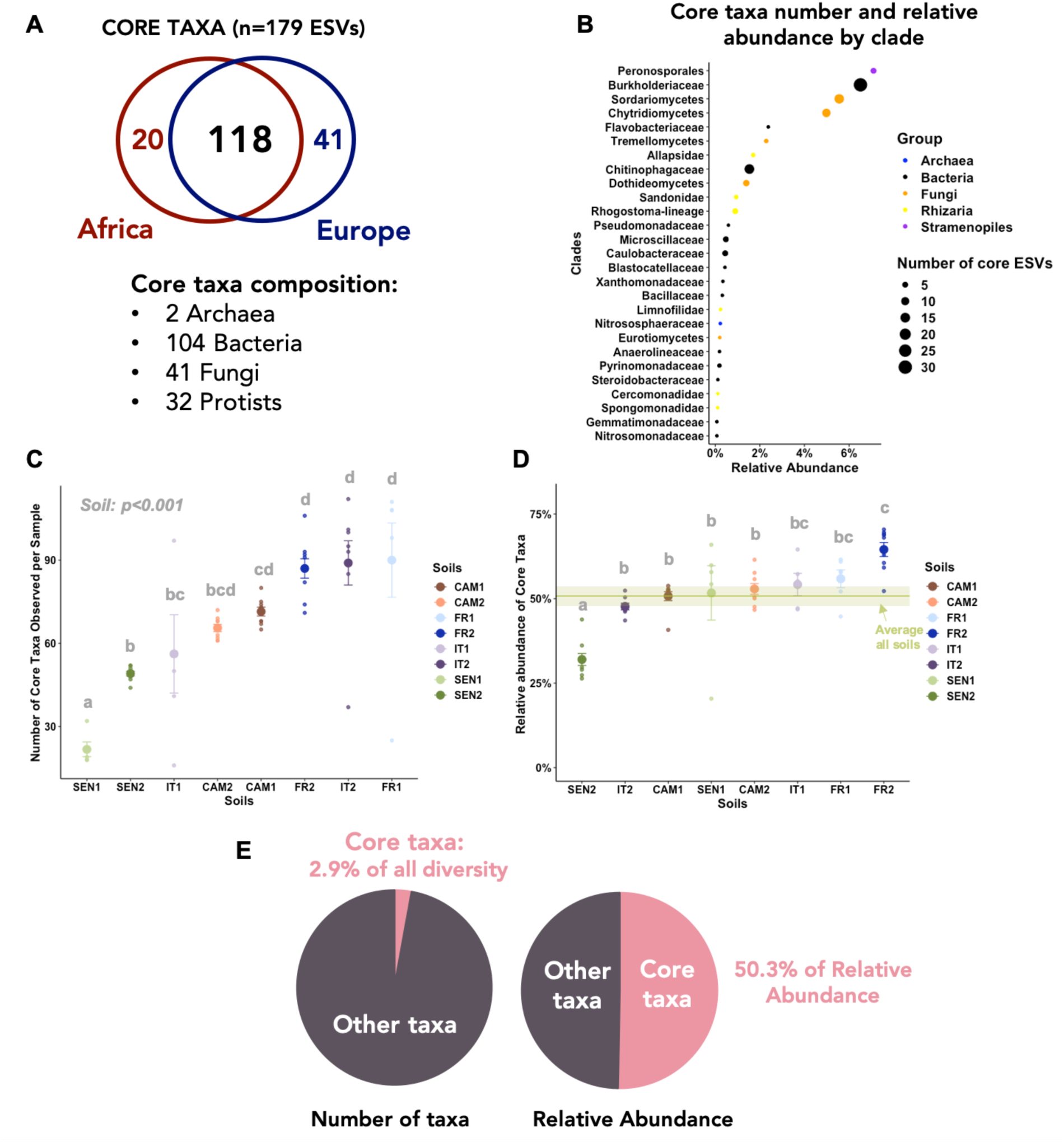
A) Venn diagram showing that among the 179 taxa identified as core taxa, 118 taxa were present in both African and European soils and 20 taxa were specific of African soils and 41 of European soils. B) A bubble plot presenting the number of core taxa and their cumulative relative abundance by clade. C) Number and D) cumulative relative abundance of the core taxa present in each sample in the eight soils. Note that on the x axis the soils are ordered from low to high y-axis value. The green line represents the average across all samples and the associated confidence interval (95%). E) Pie charts showing that while the core taxa represent only 2.9% of the total diversity, their cumulative relative abundance is 50.3% across all samples (half of the reads).

We also studied the relative abundance of each core taxon in the different soils and found that the core taxa clustered in two main groups based on their distribution patterns across all samples (Fig. 6). These results show that even if most core taxa are present in both African and European soils, they are generally very abundant only in soils from one continent and not the other. The 98 taxa affiliated to the cluster 1 were predominantly very abundant in European soils and the 81 ESVs of the cluster 2 were most abundant in African soils. Interestingly, the proportion of prokaryotes and eukaryotes were different between the 2 clusters. The cluster 1 (dominant in European soils) was principally composed of bacterial taxa (65%) and fungal taxa (19%) or protists were less represented, while the cluster 2 (dominant in African soils) was dominated by eukaryotes (28% Fungi, 16% Rhizaria, 5% Stramenopiles) and presented 48% of bacterial taxa (Fig. 6).

**Fig. 6:**
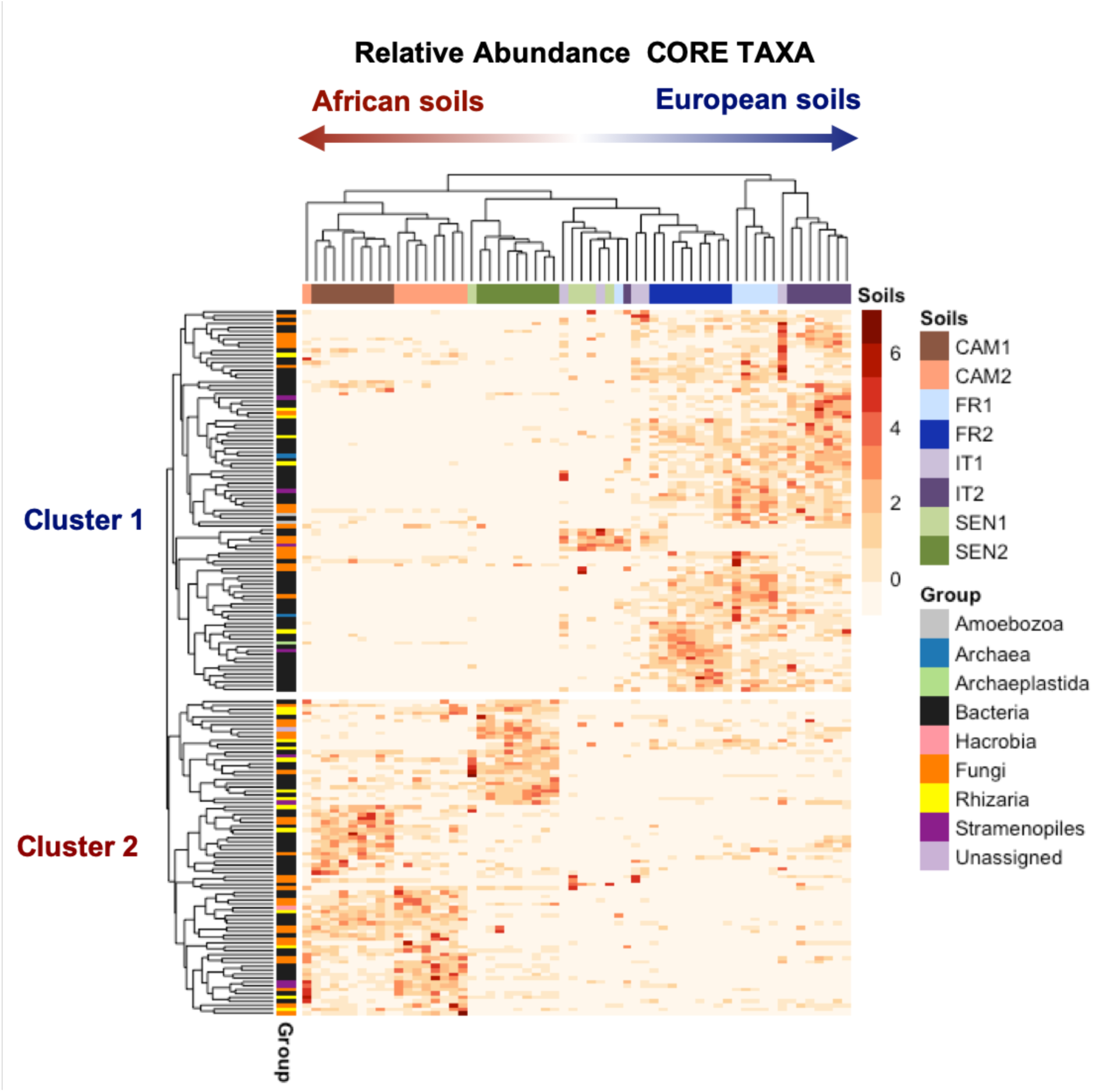
Heatmap representing the relative abundance of the 179 core taxa in the different soils. Based on the distribution of the taxa in the samples, the taxa clustered in two main groups principally associated to the continent (Africa vs Europe) where the soil was collected (but see soil SENb2). The cluster 1 regroups 98 ESVs (65% Bacteria, 19% Fungi) and the cluster 2 regroups 81 ESVs (48% Bacteria, 28% Fungi, 16% Rhizaria).

### Co-occurrence network of the core wheat microbiome and identification of hub taxa

The cross-domain network of the 179 core taxa computed with the SParse InversE Covariance estimation for Ecological Association Inference (SPIEC-EASI) was comprised of two main components, one being dominated by bacterial taxa and the other dominated by fungal and cercozoan (protists) taxa (Fig. 7A). The two main components broadly corresponded to the two clusters identified earlier based on the taxa relative abundance across all samples, with the cluster 1 gathering taxa mainly abundant in European soils and the cluster 2 in African soils (Fig. 7B). In the network, the predicted interactions were predominantly positive (440 positive edges vs 56 negative edges) and for each ESV, the average number of associations with other taxa was 5.54 (node degree). Eight taxa were identified as potential “hub” taxa based on their centrality and number of associations in the network. These hub taxa are hypothesized to be keystone taxa or key connector taxa in a community because of their central position in the network and their high connectivity. Three fungal ESVs that were extremely prevalent across all soils (60 to 85% of samples) were identified as hub taxa: *Mortierella* (Mucoromycota), *Exophiala* (Ascomycota) and an uncultured Chytridomycetes (Chytridiomycota). The other five hub taxa were bacterial ESVs in the Gammaproteobacteria phylum, one ESVs affiliated to the genus Thermomonas and two ESVs from the Burkholderiaceae family: *Massilia* and an uncultured taxon; in the Deltaproteobacteria class, one ESVs affiliated to the genus *Anaeromyxobacter*; in the class Bacteroidia, one ESV affiliated to the species *Niastella koreensis* and finally an ESV from the class Blastocatellia from an uncultured Blastocatellaceae. Interestingly, five of these potential hub taxa were connectors between the two main components of the network described above, with these connections being often negative edges.

**Fig. 7:**
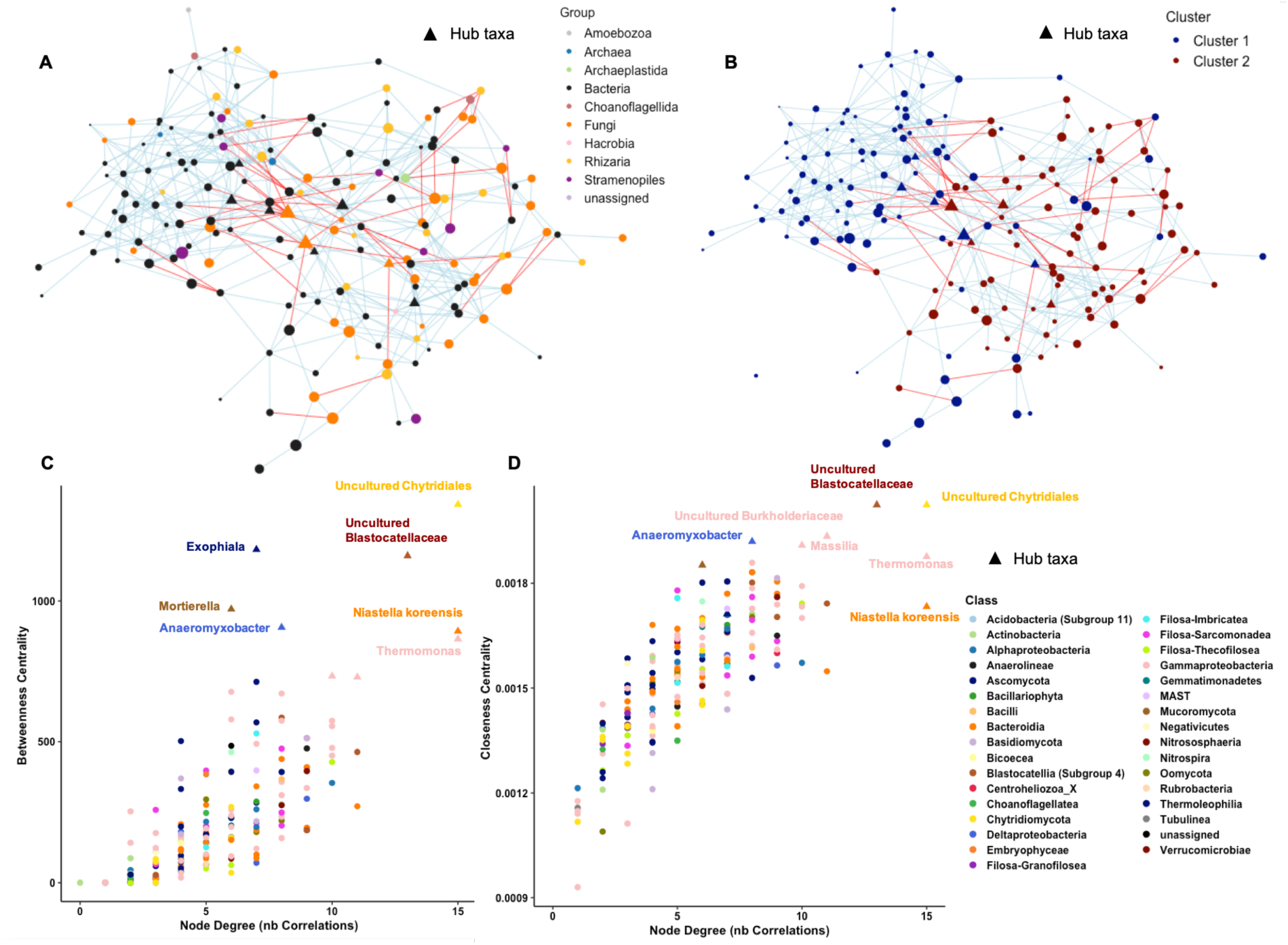
Cross domain network of the rhizosphere core taxa including both prokaryotes and eukaryotes (179 ESVs, n=60 samples). The lines (edges) between the taxa (nodes) represent the predicted interactions either positive (light blue) or negative (red). The same network is presented twice but the nodes representing the different taxa are colored by their domain or eukaryotic supergroup in the left figure (A) and are colored by their cluster affiliation in the right figure (B) as defined in the heatmap in Fig. 5. The core taxa affiliated to the cluster 1 generally had higher relative abundance in European soils, while the cluster 2 taxa were most abundant in African soils. The “Hub” taxa represented with a triangle symbol in the networks were identified as the ones presenting the highest centrality (betweenness and closeness centrality) and connectivity (node degree) in the network (panels C and D). These hubs are potential keystone taxa or key connectors in the community. The taxonomic affiliations of the hub taxa are indicated in the bottom plots and they are highlighted with triangle symbols.

A list of the most prevalent core taxa across all samples (>40% of samples) and of the ones identified as hub taxa are presented in Table 2 (complete list in Additional file 1 with sequence and taxonomy information). This table also indicates the taxa that were previously reported in the literature as core or hub taxa in the wheat microbiome. The most prevalent taxa across all samples were three eukaryotic ESVs affiliated to the fungal genera *Mortierella* (85% of samples) and *Fusarium* (82%) and a cercozoan zooflagellate belonging to the Allapsidae family (77%). The most prevalent bacterial taxa were *Bradyrhizobium japonicum* (72% of samples) and an *Arthrobacter* (70%). We observed that the relative abundance across all samples of the core taxa varied greatly (min=0.02%, max=5.7%) and that a majority of the core taxa were relatively rare in terms of abundance in the community (median=0.1%). This was also true for the eight taxa identified as hubs for which only two fungal taxa had a high relative abundance (4.24% and 1.71%) and the six others had relative abundance inferior to 0.3%. Using a complementary dataset not presented here, we determined that at least 33% of the core taxa were also observed in the wheat endosphere in the same plant samples (Table 2).

**Table 2:** Summary information on the most prevalent core taxa (>40% of samples) and hub taxa. The “Other Report” column indicates published references that reported these taxa as hub or core taxa in the wheat microbiome.

| Group | Phylum/Class | Family | Final affiliation | Preval | Rel Abund | Hub Taxa | Present in Endo | Cluster | Other Reports |
| --- | --- | --- | --- | --- | --- | --- | --- | --- | --- |
| Fungi | Mucoromycota | Mucoromycotina | Mortierella | 85% | 4.24% | Hub | Endo | 1 | Ref 1, 2, 3 |
| Fungi | Ascomycota | Eurotiomycetes | Exophiala | 62% | 0.16% | Hub |  | 1 | Ref 2 |
| Fungi | Chytridiomycota | Chytridiomycetes | Uncultured Chytridiomycetes | 60% | 1.71% | Hub | Endo | 2 |  |
| Bacteria | Gammaproteobacteria | Burkholderiaceae | Massilia | 47% | 0.25% | Hub | Endo | 2 | Ref 5 |
| Bacteria | Blastocatellia | Blastocatellaceae | Uncultured Blastocatellaceae | 43% | 0.30% | Hub | Endo | 1 |  |
| Bacteria | Gammaproteobacteria | Burkholderiaceae | Uncultured Burkholderiaceae | 42% | 0.20% | Hub |  | 1 |  |
| Bacteria | Deltaproteobacteria | Archangiaceae | Anaeromyxobacter | 38% | 0.05% | Hub |  | 2 |  |
| Bacteria | Gammaproteobacteria | Xanthomonadaceae | Thermomonas | 38% | 0.25% | Hub |  | 1 |  |
| Bacteria | Bacteroidia | Chitinophagaceae | Niastella korensis | 32% | 0.09% | Hub | Endo | 2 |  |
| Fungi | Ascomycota | Sordariomycetes | Fusarium | 82% | 1.69% |  | Endo | 1 | Ref 2 |
| Rhizaria | Filosa-Sarcomonadea | Allapsidae | Allapsidae (Group-Te) | 77% | 0.91% |  |  | 2 |  |
| Bacteria | Alphaproteobacteria | Xanthobacteraceae | Bradyrhizobium japonicum | 72% | 0.14% |  | Endo | 2 |  |
| Bacteria | Actinobacteria | Micrococcaceae | Arthrobacter | 70% | 0.19% |  |  | 2 |  |
| Rhizaria | Filosa-Granofilosea | Limnophiliidae | Limnophila borokensis | 70% | 0.20% |  |  | 2 |  |
| Bacteria | Nitrospira | Nitrospiraceae | Nitrospira | 67% | 0.14% |  |  | 1 |  |
| Bacteria | Gammaproteobacteria | Enterobacteriaceae | Pantoea agglomerans | 67% | 0.39% |  | Endo | 2 |  |
| Rhizaria | Filosa-Sarcomonadea | Paracercomonadidae | Paracercomonas | 63% | 0.15% |  | Endo | 1 |  |
| Bacteria | Gammaproteobacteria | Burkholderiaceae | Paucibacter | 62% | 0.09% |  |  | 2 |  |
| Fungi | Ascomycota | Sordariomycetes | Chaetomium | 60% | 0.59% |  | Endo | 1 | Ref 2, 4 |
| Bacteria | Gammaproteobacteria | Burkholderiaceae | Ramlibacter | 60% | 0.09% |  |  | 2 |  |
| Bacteria | Actinobacteria | Geodermatophilaceae | Blastococcus | 57% | 0.13% |  |  | 1 |  |
| Bacteria | Gammaproteobacteria | Burkholderiaceae | Noviherbaspirillum | 57% | 0.21% |  |  | 2 |  |
| Bacteria | Alphaproteobacteria | Caulobacteraceae | Phenylobacterium | 57% | 0.10% |  |  | 2 |  |
| Fungi | Ascomycota | Sordariomycetes | Bionectria | 55% | 0.31% |  |  | 2 |  |
| Fungi | Ascomycota | Dothideomycetes | Uncultured Dothideomycetes | 55% | 0.31% |  | Endo | 1 |  |
| Rhizaria | Filosa-Sarcomonadea | Cercomonadidae | Eocercomonas | 53% | 0.05% |  |  | 2 |  |
| Bacteria | Bacteroidia | Chitinophagaceae | Flavitalea | 50% | 0.08% |  |  | 2 |  |
| Fungi | Ascomycota | Sordariomycetes | Uncultured Sordariomycetes | 50% | 0.25% |  | Endo | 2 |  |
| Bacteria | Gammaproteobacteria | Burkholderiaceae | Massilia | 48% | 0.31% |  | Endo | 1 |  |
| Bacteria | Bacteroidia | Chitinophagaceae | Uncultured Chitinophagaceae | 48% | 0.42% |  | Endo | 1 |  |
| Fungi | Ascomycota | Dothideomycetes | Uncultured Pleosporales | 48% | 0.18% |  | Endo | 2 |  |
| Bacteria | Gammaproteobacteria | Burkholderiaceae | Ramlibacter | 47% | 0.08% |  |  | 2 |  |
| Rhizaria | Filosa-Imbricatea | Thaumatomonadidae | Thaumatomonas | 47% | 0.22% |  |  | 2 |  |
| Fungi | Ascomycota | Dothideomycetes | Uncultured Dothideomycetes | 47% | 0.22% |  |  | 2 |  |
| Fungi | Chytridiomycota | Chytridiomycetes | Spizellomyces-Rhizophlyctidales | 45% | 0.45% |  |  | 2 |  |
| Fungi | Chytridiomycota | Chytridiomycetes | Spizellomyces-Rhizophlyctidales | 45% | 0.19% |  |  | 2 |  |
| Fungi | Ascomycota | Sordariomycetes | Uncultured Sordariomycetes | 45% | 0.37% |  |  | 1 |  |
| Bacteria | Alphaproteobacteria | Caulobacteraceae | Asticcacaulis | 43% | 0.12% |  |  | 1 |  |
| Fungi | Basidiomycota | Tremellomycetes | Cryptococcus | 43% | 1.40% |  | Endo | 1 |  |
| Stramenopile | Oomycota | Peronosporales | Pythium | 43% | 0.46% |  | Endo | 2 |  |
| Fungi | Chytridiomycota | Chytridiomycetes | Rhizophlyctis | 43% | 0.25% |  |  | 1 |  |
| Bacteria | Gammaproteobacteria | Burkholderiaceae | Uncultured Burkholderiaceae | 43% | 0.39% |  |  | 1 |  |
| Bacteria | Bacteroidia | Microscillaceae | Uncultured Microscillaceae | 43% | 0.19% |  |  | 1 |  |
| Bacteria | Gammaproteobacteria | Burkholderiaceae | Acidovorax | 42% | 0.79% |  | Endo | 1 |  |
| Rhizaria | Filosa-Sarcomonadea | Allapsidae | Allapsidae (Group-Te) | 42% | 0.66% |  |  | 2 |  |
| Bacteria | Bacilli | Bacillaceae | Bacillus | 42% | 0.23% |  |  | 2 |  |
| Bacteria | Bacteroidia | Chitinophagaceae | Flavisolibacter | 42% | 0.07% |  |  | 2 |  |
| Bacteria | Bacteroidia | Chitinophagaceae | Flavisolibacter | 42% | 0.11% |  |  | 2 |  |
| Bacteria | Gammaproteobacteria | Methylophilaceae | Methylophilera | 42% | 0.04% |  |  | 2 |  |
| Bacteria | Alphaproteobacteria | Sphingomonadaceae | Sphingomonas | 42% | 0.07% |  |  | 2 |  |
| Fungi | Chytridiomycota | Chytridiomycetes | Uncultured Chytridiomycetes | 42% | 0.76% |  |  | 2 |  |
| Stramenopile | MAST | MAST-12C | Uncultured MAST-12C | 42% | 0.08% |  |  | 2 |  |
| Rhizaria | Filosa-Sarcomonadea | Sandonidae | Uncultured Sandonidae | 42% | 0.81% |  | Endo | 1 |  |
| Fungi | Ascomycota | Sordariomycetes | Uncultured Sordariomycetes | 42% | 0.13% |  | Endo | 2 |  |

Preval= Prevalence; Rel Abund = Relative Abundance; Present in Endo = Present in Endosphere
Ref 1: Hartman et al. 2018; Ref 2: Schlatter et al. 2018; Ref 3: Borrell et al. 2017 Ref 4 : Fan et al. 2018, Ref 5 : Granzow et al. 2017

## Discussion

In this study, we took an integrative approach to characterize the rhizosphere microbiome (i.e. soil tightly bound to the roots) of multiple wheat genotypes by considering the total microbial diversity, including archaea, bacteria, fungi and protists in soils from different countries. We show that the main drivers of the wheat rhizosphere microbiome are the culture soil and agricultural practices, while wheat genotypes had very limited effects. Across eight contrasted soils from two continents and three wheat genotypes, we found that 179 prokaryotic and eukaryotic taxa were consistently associated to the wheat rhizosphere, constituting a core microbiome. In addition to being prevalent, these few core taxa were highly abundant and collectively represented 50% of the relative abundance of the wheat microbiome on average. This work enabled the identification of a taxa short-list for future wheat microbiome research that will be targets for culturomic, genomic and synthetic community studies to develop microbiome engineering in agriculture.

### The wheat rhizosphere microbiome is shaped by the soil and agricultural practices but not by the plant genotype

By growing eight genotypes in the same soil, we observed that wheat varieties had very limited effects on rhizosphere microbiome diversity (richness and phylogenetic diversity) and structure. The variability among replicates of a same genotype was often larger than between genotypes. These results were confirmed in our second sub-experiment where we did not find a significant effect of the three genotypes selected in the eight test soils (only 2% of variance explained). These results are consistent with previous work showing a small effect of the wheat genotype explaining only 1 to 4% of the total variance in the community or effects appearing only after two years of cultivation on the field (Donn *et al.* 2015; Corneo *et al.* 2016; Mahoney, Yin and Hulbert 2017). Hence, the lack of genotype effect in our study could be explained by the short duration of our experiment with the harvest taking place at the vegetative stage when root microbiome is not fully “mature” (Donn *et al.* 2015; Gdanetz and Trail 2017).

Consistent with the literature, the main factor shaping the wheat rhizosphere microbiome diversity and structure was the test soil (57% of the variance). A clear separation between African and European soils was observed indicating potential continental effects on plant microbiome structure. It should be noted that the two European soils were closely clustered on the NMDS, while the Cameroonian and Senegalese soils were clearly separated on the second axis. We also observed distinct community structure between the two soils sampled from each country, that were the strongest for the two Senegalese soils. These separations could be explained by differences in soil parameters, especially pH (lower pH in African soils in our study) that has been found to be the main soil parameter influencing wheat microbiomes at a regional scale in China (Fan *et al.* 2017).

Large differences in taxa richness and phylogenetic diversity were present between the eight soils (100 to 380 total taxa), indicating that not only the microbiome composition was different but also the number of taxa selected by the plant, as already shown in the wheat rhizosphere (Corneo *et al.* 2016; Mahoney, Yin and Hulbert 2017; Yin *et al.* 2017). We did not observe clear diversity patterns associated to countries, for instance the two Senegalese soils had respectively the lowest and highest phylogenetic diversity of the study and large differences in ESV richness were also observed between the two Italian soils. In contrast, the diversity indices of the two Cameroonian or French soils were not statistically different from each other despite large differences in soil properties and agricultural practices. Interestingly across all soils, prokaryotic diversity was positively correlated to eukaryotic diversity signifying that the environmental conditions favourable to high diversity are similar across all microorganisms. These results show in the contrasted eight soils studied that prokaryotic diversity is 2 to 3 times higher than eukaryotic diversity in the wheat rhizosphere.

Agricultural practices were also a significant driver of microbiome structure (10% of the variance) in Italian and French soils in which we compared rhizosphere microbiomes between conventional and organic farming. However, the taxa richness was significantly affected by agricultural practices only in the Italian soils. These results are very similar to the findings of Hartman et al. (2018)(Hartman *et al.* 2018) that observed more effects of conventional vs organic practices on wheat microbiome structure (9-10% of the variance) than on diversity but also with other studies in various cropping systems (Gdanetz and Trail 2017; Granzow *et al.* 2017). The IT2 soil sampled from an organic farming field presented a higher taxa richness (56% higher) than IT1. It is noteworthy that this IT1 soil has been sampled from a field uninterruptedly sowed for 15 years with small grain cereals (wheat and barley) that can be considered a long-term monoculture that were cultivated adopting conventional farming procedures. The lower microbial diversity of IT1 soil in comparison with the other European soils can be explained by the different hypotheses. First, the higher diversity in the IT2 organic field could be associated to the addition of new microorganisms with amendments in organic farming. Moreover, crop rotation compared to long-term monocropping is leading on average to a ∼15.1% increase in microbial richness (Venter, Jacobs and Hawkins 2016).

The rhizosphere microbiome was dominated by the bacterial family Burkholderiaceae (10% of all sequences), oomycetes from the order Peronosporales (8.6%) and fungi from the Sordariomycetes (7.3%) and Chytridiomycetes (6.5%) classes. The diversity of protists, especially Cercozoans, was extremely high, representing half of the eukaryotic diversity in the microbiome, with many abundant predator taxa affiliated to *Rhogostoma* (testate amoebae), *Allapsidae-Group Te* (glissomonads) and *Sandonidae* (glissomonads). This study shows that non-fungal eukaryotes are diverse and abundant in the wheat rhizosphere and they deserve more attention to determine their functional roles in nutrient cycling and in controlling microbiome structure through microbial predation.

While many taxa were found to be shared between different genotypes and soils (see core microbiome section below), we also observed microbial taxa that were specific of some soils or agricultural practices and that were responsible for the significant shifts in community structure and diversity presented above. In particular, ESVs affiliated to the Acidobacteria, Chloroflexi and Bacteroidetes phyla were often found only in African or European soils but not in both, suggesting a continental phylogenetic signal or full clades sensitive to differences in soil pH. Differentially abundant taxa between conventional and organic farming were identified in almost all groups (bacteria, oomycetes, fungi, cercozoans) indicating a clear restructuring of the microbiome as a whole. Interestingly, potential plant pathogenic taxa were found to be enriched in conventional (*Phytophtora*) or organic farming (*Pythium*).

Using an integrative approach of the plant microbiome, contrary to most studies that separate bacteria and fungi in their analyses, we intentionally grouped all taxa in our structure and diversity analyses together to present a global view of the wheat microbiome as it occurs in nature. With this approach, we show that the soil and agricultural practices are the key drivers of wheat rhizosphere microbiome and that a large part its micro-eukaryotic diversity is constituted of protists that has been so far overlooked in the plant holobiont.

### Evidences for a wheat core microbiome and identification of hub taxa

Many prokaryotic and eukaryotic taxa were found to be consistently associated to wheat rhizospheres from multiple genotypes and soils, with the exact same 16S or 18S rRNA sequence variants detected in soils sampled thousands of kilometers apart. More specifically, we identified 179 taxa that constitute the core microbiome of the wheat rhizosphere in eight African and European soils. These core taxa where highly prevalent (present in multiple soils) but also very abundant which was not necessarily expected as we did not impose a relative abundance threshold to identify core taxa. Collectively, these 179 taxa (2.9% of the total diversity) represented 50% of the reads in the dataset, suggesting an important biomass of these microorganisms in the rhizosphere. These findings show that under very contrasted conditions (continents, soil types, farming practices, wheat genotypes), wheat associates with the same microbial species or strains (ESV level) which indicate a potential co-evolution between wheat and these microorganisms. These results are consistent with large-scale microbiome studies that identified the core microbiome of different plants like *Arabidopsis thaliana* (Lundberg *et al.* 2012) or sugarcane (Hamonts *et al.* 2018) and a global soil microbiome study that highlighted that only 500 OTUs account for half of soil microbial communities worldwide (Delgado-Baquerizo *et al.* 2018).

The wheat rhizosphere microbiome can be diverse and variable (100 to 380 taxa) but these findings show that it can be decomposed in two parts: a core microbiome presenting a low diversity that is stable and abundant across conditions; and an “accessory” microbiome that is extremely diverse and condition-specific (Vandenkoornhuyse *et al.* 2015). This observation is consistent with a previous wheat microbiome study that showed that highly co-occurring taxa (i.e. potential core taxa) were not affected by cropping practices and that the “accessory” wheat microbiome could be manipulated by changes in farming practices (Hartman *et al.* 2018). The high prevalence and abundance of the core taxa identified suggest an ecological significance in the root habitat and warrants further research to isolate and phenotype these organisms that are mainly uncultured (i.e. 62 core taxa without genus affiliations). Based only on amplicon sequencing data, we cannot determine the role of these microorganisms for plant fitness or if the absence of some of these core taxa could impact the wheat microbiome or plant health. Thus, here we provide a taxa list composed of 2 archaea, 103 bacteria, 41 fungi and 32 protists that should be the target of future culturomics, metagenomics and for the creation of wheat synthetic microbiomes.

Most of these core taxa were present in both African and European soils (118 taxa) but based on their relative abundance they grouped into two clusters, with taxa having a higher abundance in either European soils (Cluster 1) or African soils (Cluster 2). These results were confirmed in the cross-domain network of all core taxa that separated into two main components that broadly corresponded to the two clusters. Future synthetic community studies should consider and evaluate the effects of contrasted relative abundances of the core taxa to represent realistic structuration of the wheat microbiomes.

Among all core taxa, a fungal ESV affiliated to the genus *Mortierella* (phylum Mucoromycota) occupied a crucial place in the wheat microbiome. This taxon had the highest prevalence (85% of samples) and relative abundance in the dataset (4.2%) and was identified as a hub taxon in the cross-domain network. This *Mortierella* ESV was detected on the wheat rhizoplane but also in the endosphere in a complementary dataset not presented in this article. This fungus is described as a saprotroph-symbiotroph (Nguyen *et al.* 2016) capable of solubilizing phosphate in the rhizosphere (Zhang *et al.* 2011). Our results are consistent with other studies that identified *Mortierella* as a keystone taxa of the wheat rhizosphere unsensitive to cropping practices in Switzerland or the USA (Table 2, Hartman *et al.* 2018; Schlatter *et al.* 2018) with potential positive effects on wheat yield in Canada (Borrell *et al.* 2017). Altogether, these findings encourage future research to develop targeted cultivation approaches of the wheat-associated *Mortierella* fungi that seem to play a central role in the wheat microbiome across the world. The other most prevalent fungal core taxa were affiliated to the genera *Fusarium* (Ascomycota, 82% of samples), *Exophiala* (Ascomycota, 62% of samples, hub) and *Chaetomium* (Ascomycota, 60% of samples) described as pathotrophs or saprotrophs that also have been observed on wheat roots in different cropping systems and described as hub taxa (Gdanetz and Trail 2017; Fan *et al.* 2018; Schlatter *et al.* 2018).

The taxonomic group that exhibited the highest number of core taxa was bacteria (103 taxa) with the dominant family Burkholderiaceae presenting the highest number of core taxa (28 taxa) including two hub taxa. In particular, an ESV affiliated to the genus *Massilia* was identified as a hub taxon in the wheat microbiome of our study. This taxon is known as a copiotrophic root colonizer (Ofek, Hadar and Minz 2012) found to be dominant in the wheat and maize rhizosphere (Li *et al.* 2014; Granzow *et al.* 2017). The most prevalent bacterial core taxon was *Bradyrhizobium japonicum* (Alphaproteobacteria, 72% of samples), a species frequently found in root-associated microbiomes, and which can fix nitrogen in symbiosis with legumes (especially soybean).

Here, we show also that archaea and protists that are microbial groups generally ignored in most plant microbiome studies were characterized as wheat core taxa. The two archaea identified as core taxa were both affiliated to the genus *Nitrososphaera* (Phylum Thaumarcheota) that are involved in the nitrogen cycle as ammonia-oxidizers in the nitrification process (oxidation of ammonia in nitrate) (Tourna *et al.* 2011) and are very abundant in soils (Pester *et al.* 2012; Simonin *et al.* 2016). Interestingly, another nitrifier taxa affiliated to the bacterial genus *Nitrospira* (Nitrospira phylum, 67% of all samples) was also identified as core taxon which suggests that microorganisms involved in nitrification could play an important role for nitrogen availability on roots. *Nitrososphaera* taxa were described as hubs in the wheat rhizosphere (Fan *et al.* 2018) and as highly prevalent in the rhizosphere and rhizoplane of *Arabidopsis thaliana* and maize (Carvalhais *et al.* 2015; Walters *et al.* 2018), indicating that root surfaces represent a common habitat for these archaea.

Among protists, the most prevalent core taxa were all cercozoans from the phyla Filosa-Sarcomonadae and Filosa-Granofilosea affiliated to *Allapsidae – Group Te* (77% of samples), *Limonofila borokensis* (70%) and *Paracercomonas* (63%). These three taxa have also been identified as extremely common and abundant in the rhizosphere of *Arabidopsis thaliana* (Sapp *et al.* 2018) and *Allapsidae – Group Te* as a hub taxon in the maize rhizosphere (Zhao *et al.* 2019). Five Oomycetes were also categorized as core taxa with three of them affiliated to the *Pythium* genus and one to *Phytophthora* that are both described as potential plant pathogens. These observations are consistent with previous plant microbiome studies considering oomycetes diversity, that always described *Pythium* as the most dominant taxa among oomycetes (Durán *et al.* 2018; Hassani, Durán and Hacquard 2018; Sapp *et al.* 2018). These results confirm that protists are an integral part of the plant holobiont and their roles in controlling microbial populations through predation, disease incidence or contribution to nutrient cycles through the microbial loop deserve more attention (Gao *et al.* 2019).

In conclusion, this work presents a detailed characterization of the wheat rhizosphere microbiome under contrasted environmental conditions and details a list of microorganisms identified as core and hub taxa. In this list of 179 prokaryotic and eukaryotic taxa, 62 are uncultured taxa (no genus affiliation) and only 11 taxa have a species affiliation, suggesting that almost nothing is known about the ecology of the key microorganisms associated to a major crop like wheat. Future research efforts are needed to try to cultivate these microorganisms associated to wheat as they could unlock new knowledge and biotechnological resources to improve crop yields and resistance to diseases in a sustainable way. These efforts should be conducted with an integrative approach of the plant holobiont by not only focusing on bacteria and fungi but also considering archaea, oomycetes and cercozoans.

## Additional files

**Additional file 1**: An XLSX table containing a tab with the full list of the 179 core taxa with taxonomic, prevalence, relative abundance and sequence information. A second tab presents the list of taxa found to be associated to 7-8 wheat genotypes in the FR2 soil and the third tab provides the list of taxa significantly affected by farming practices.

## Acknowledgements

We thank Dr. Jacques Le Gouis from INRA Clermont-Ferrand (UMR GDEC) for help in sampling French soils and for providing several wheat seeds cultivars. We thank Dr Honore Tekeu, M. Yves Tchietchoua and Ms Adrienne Ngo Ngom for soil sampling at Tubah area in Cameroon and Institute of Agricultural Research for Development (IRAD) that hosted all activities of the wheat research project in Cameroon. We also thank Dr. Fatou NDOYE and Ms Achaitou DJIBO WAZIRI for Senegalese soils sampling. For lab assistance, we thank Isabelle Rimbault and Nathalia Arias Rojas.

## Supplementary Information

**Fig S1:**
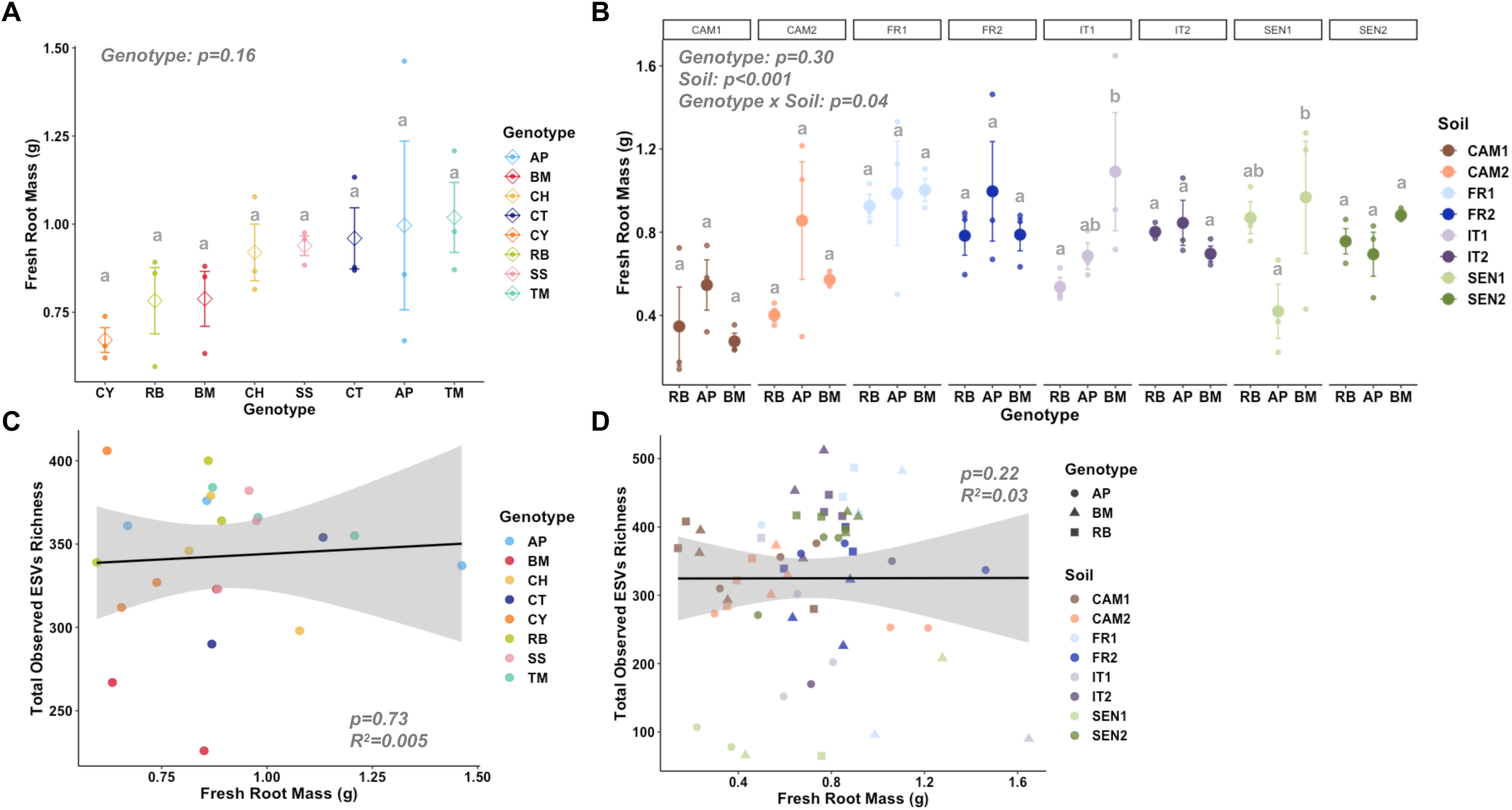
Results of the Sub-experiment 1 & 2: Fresh root mass measurements for A) the eight genotypes grown in FR2 soil (sub-experiment 1) and B) the three genotypes grown in the eight different soils (sub-experiment 2). Different letters represent statistical differences between the different genotypes of a same soil. The panel C and D show the lack of correlation between the fresh root mass and the total observed ESVs richness (Prokaryote + Eukaryote) for the sub-experiment 1 (FR2 soil only) and the sub-experiment 2 (eight soils).

**Fig S2:**
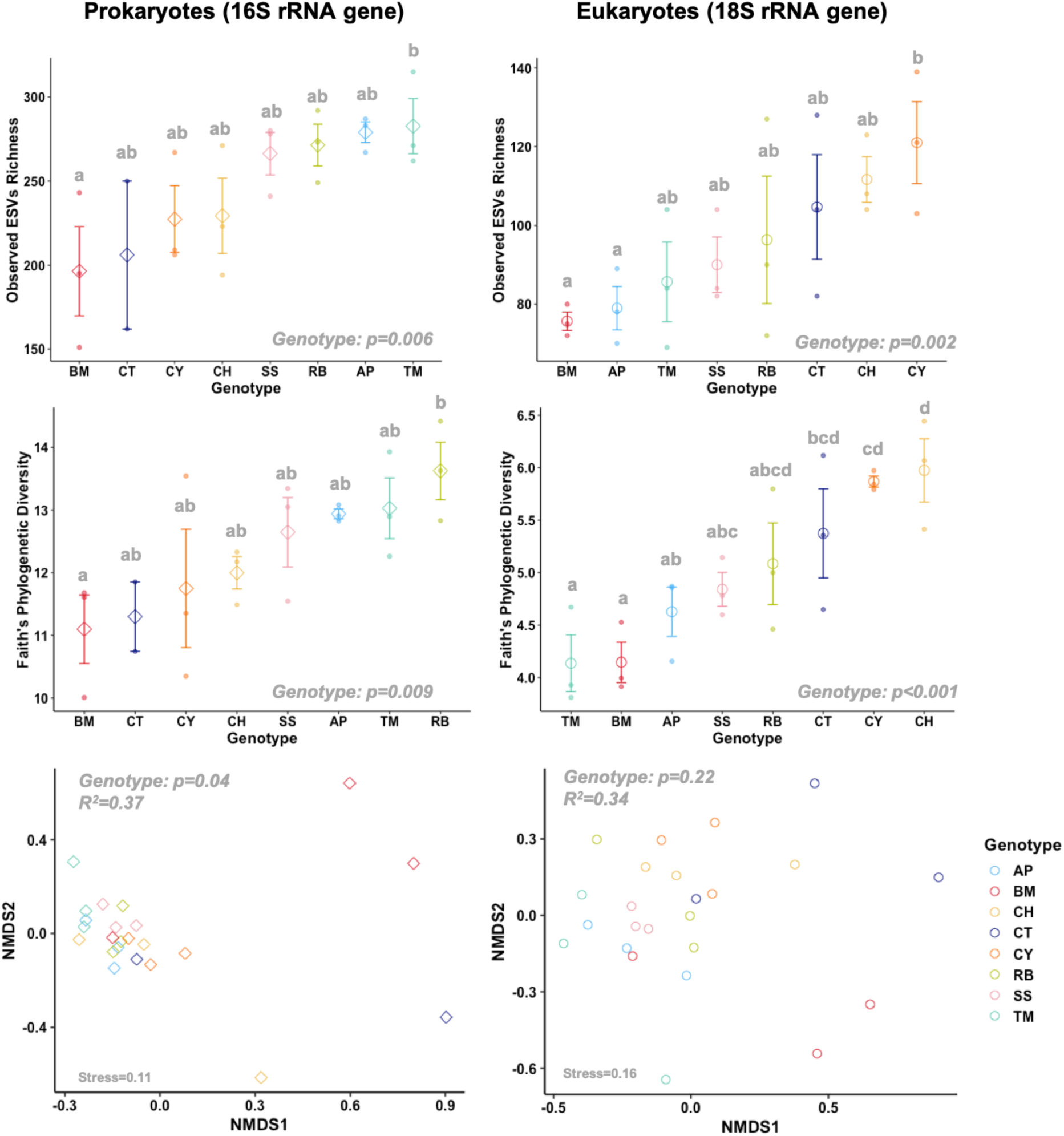
Results of the Sub-experiment 1: Effects of the wheat genotypes on the prokaryotic and eukaryotic microbial communities. Top: Observed ESV richness, Middle: Faith’s phylogenetic diversity, Bottom: Community Structure represented in a NMDS. Different letters represent statistical differences between the eight different genotypes.

**Fig S3:**
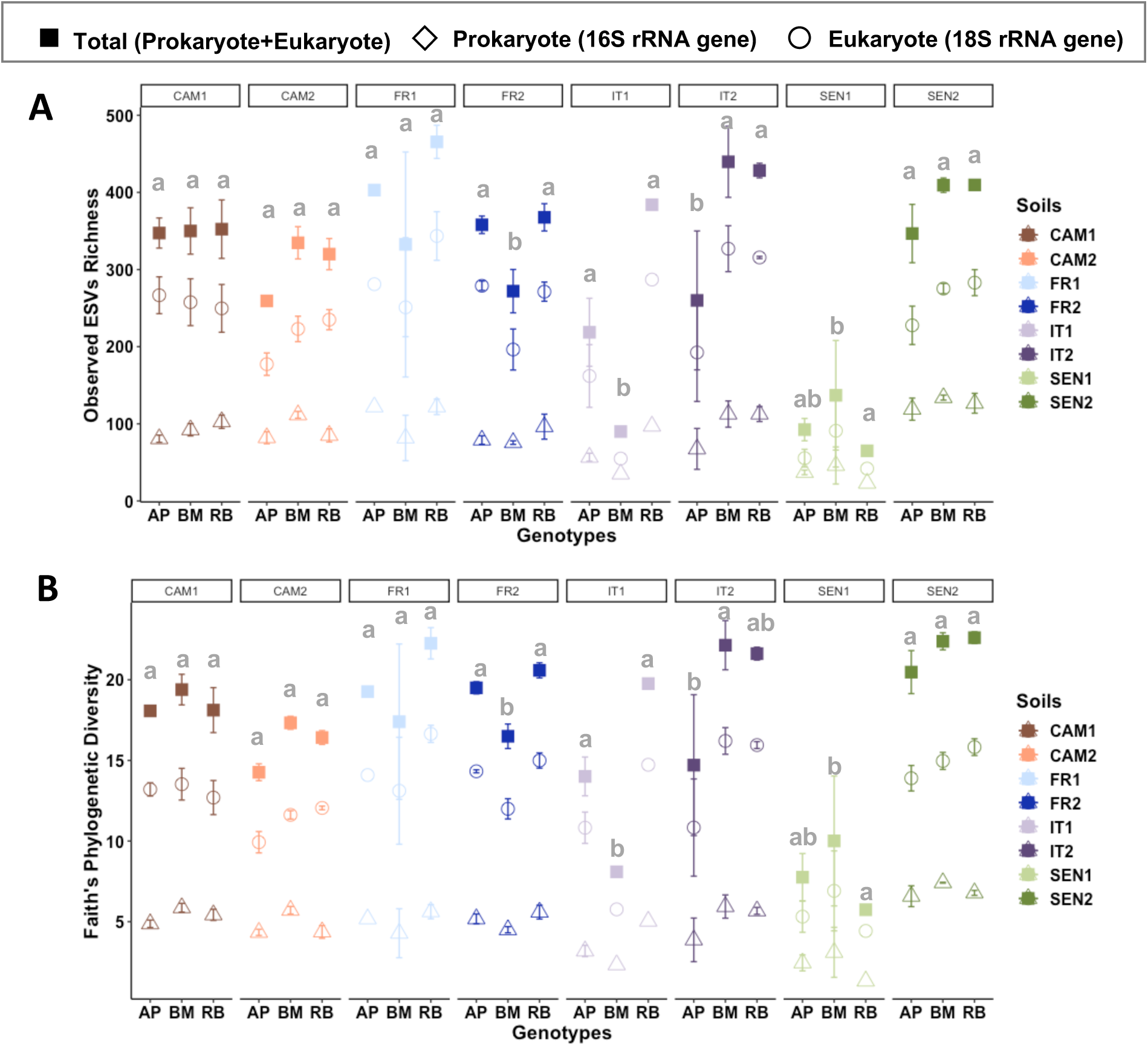
Results of the Sub-experiment 2: Effects of the soil and wheat genotypes on total, prokaryotic and eukaryotic microbial diversity. A: Observed ESV richness, B: Faith’s phylogenetic diversity. Different letters represent statistical differences for each soil in total diversity (Prokaryote + Eukaryote) between genotypes.

**Fig S4:**
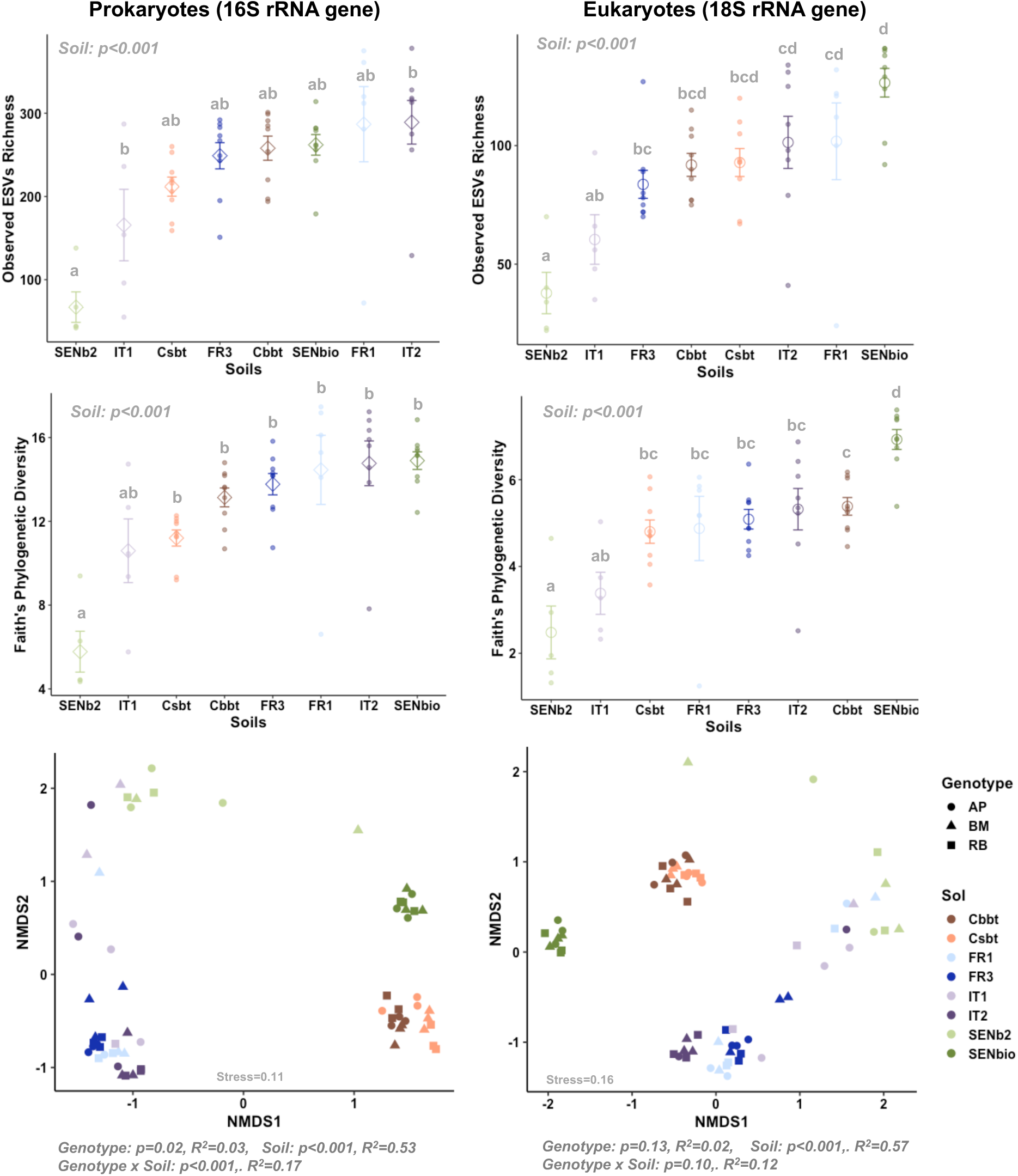
Results of the Sub-experiment 2: Effects of the soil and wheat genotypes on the prokaryotic and eukaryotic microbial communities. Top: Observed ESV richness, Middle: Faith’s phylogenetic diversity, Bottom: Community Structure represented in a NMDS. For the Observed Richness and Faith’s Phylogenetic diversity, the average and standard deviation are presented for the three genotypes combined for each soil. Different letters represent statistical differences between soils.

**Fig S5:**
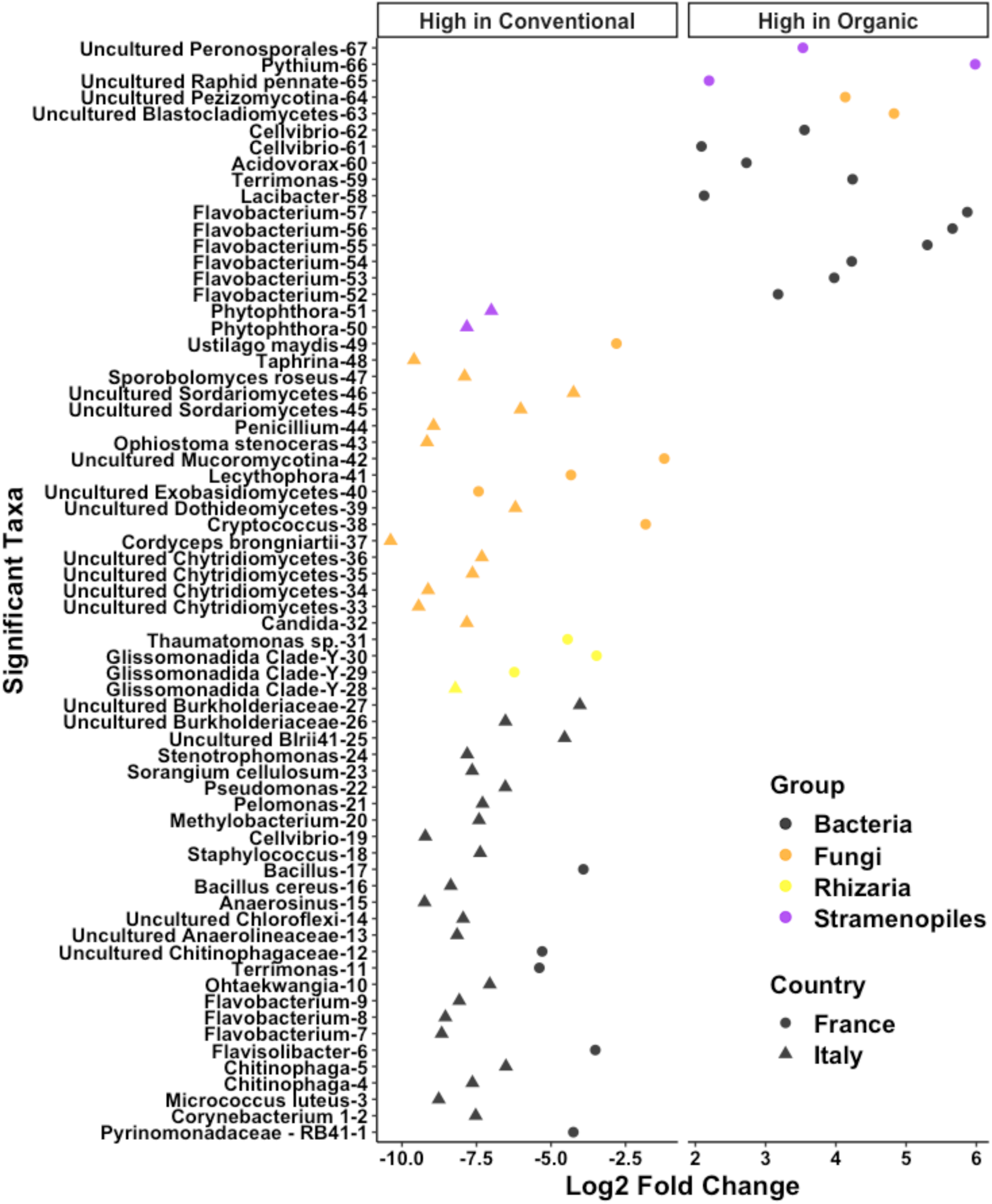
Taxa significantly enriched in Conventional or Organic agricultural practices in the French and Italian soil. Each taxa Log2 fold change relative to the other agricultural condition is indicated on the x-axis.

**Fig S6:**
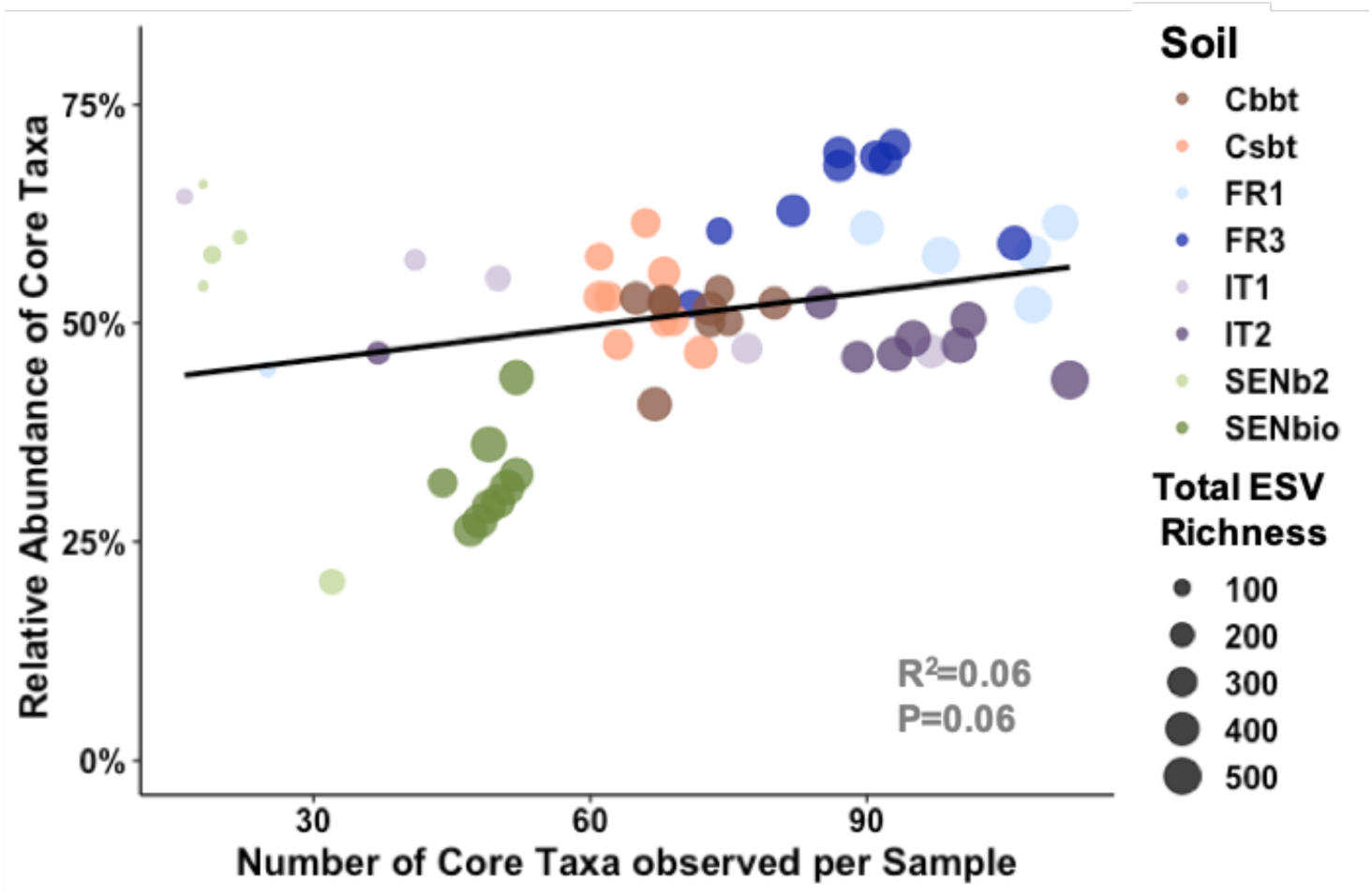
Correlation between the number of core taxa and the cumulative relative abundance of the core taxa per sample in the eight soils. The size of the points indicates the total ESV richness in the sample (prokaryotes + eukaryotes).

